# A multi-omics analysis identifies molecular features associated with heifer fertility in a case-control design including Angus and Holstein cattle

**DOI:** 10.1101/2022.12.13.520240

**Authors:** Mackenzie A. Marrella, Fernando H. Biase

**Affiliations:** School of Animal Sciences, Virginia Polytechnic Institute and State University, Blacksburg, Virginia, USA

**Keywords:** fertility, heifer, beef, dairy, transcriptome, Angus, Holstein, SNP

## Abstract

**Background:** Infertility or subfertility is a critical barrier to sustainable cattle production, including in heifers. The development of heifers that do not produce a calf within an optimum window of time is a critical factor for the profitability and sustainability of the cattle industry. The early identification of heifers with optimum fertility using molecular phenotyping is a promising approach to improving sustainability in beef and dairy cattle production.

**Results:** Using a high-density single nucleotide polymorphism (SNP) chip, we collected genotypic data from 575,053 SNPs. We also produced quantitative transcriptome data for 12,445 genes (12,105 protein-coding genes, 228 long non-coding RNAs, and 112 pseudogenes) and proteome data for 213 proteins. We identified two SNPs significantly associated with heifer fertility (rs110918927, chr12: 85648422, P = 6.7×10^-7^; and rs109366560, chr11:37666527, P = 2.6×10^-5^). We identified two genes with differential transcript abundance (eFDR ≤ 0.002) between the two groups (Fertile and Sub-Fertile): Adipocyte Plasma Membrane Associated Protein (*APMAP*, 1.16 greater abundance in the Fertile group) and Dynein Axonemal Intermediate Chain 7 (*DNAI7*, 1.23 greater abundance in the Sub-Fertile group). Our analysis revealed that the protein Alpha-ketoglutarate-dependent dioxygenase FTO was more abundant in the plasma collected from Fertile heifers relative to their Sub-Fertile counterparts (FDR < 0.05). Interestingly, two proteins did not reach the significance threshold in the model accounting for all samples (Apolipoprotein C-II, APOC2 (FDR_glmm_ = 0.06) and Lymphocyte cytosolic protein 1, LCP1 (FDR_glmm_ = 0.06)), but both proteins were less abundant in the plasma of Fertile Holstein heifers (P < 0.05). Lastly, an integrative analysis of the three datasets identified a series of features (SNPs, gene transcripts, and proteins) that can be useful for the discrimination of heifers based on their fertility. When all features were utilized together, 21 out of 22 heifers were classified correctly based on their fertility category.

**Conclusions:** Our multi-omics analyses confirm the complex nature of female fertility. Very importantly, our results also highlight differences in the molecular profile of heifers associated with fertility that transcend the constraints of breed-specific genetic background.

## BACKGROUND

The latest data from the Food and Agriculture Organization show that in 2013 more than 39% of the daily protein supply in the world was from animal-based foods (FAO-STATS). Bovine meat and milk accounted for 14.5% of the total protein supply in the world in 2013 (FAO-STATS). These numbers underscore the importance of cattle production to sustain a growing demand for protein globaly [1]. Infertility or subfertility is a critical barrier to sustainable cattle production [2], including in heifers. For example, approximately 15% [3] and 5% [4] of beef and dairy heifers, respectively, do not calve at 24 months of age. Heifers that calve at an optimum age have greater productivity and longevity in the herd [5–10]. Therefore, identifying heifers with optimum fertility is a promising approach to improve sustainability in cattle production.

The calculation of breeding values for fertility traits in heifers serves as an important tool for improving fertility overall. However, the heritabilities of breeding values for heifer fertility is often low for beef [11–17] and dairy [18–23] heifers, which contribute to a low rate of genetic change for such traits. Another potential avenue for the identification of infertile heifers is the use of molecular phenotyping [24]. The pioneering efforts focused on genome-wide association studies (GWAS) to identify genetic markers associated with heifer fertility [4, 12, 25–38], but only a few seem to be reproducible across populations [37]. More recent efforts have also focused on transcriptome [39–41] and metabolome [42] datasets characterizing these molecules in blood samples. Again, limited genes have been identified with differential transcript abundance across datasets [39]. Although promising, much research is needed for the identification of biomarkers that can help guide early management actions on infertile or sub-fertile heifers.

We carried out an experiment to test the hypothesis that differences in genetic variants, gene transcript, and protein abundance due to fertility fitness would be shared between heifers of different breeds. Our objective was to contrast genetic variants, gene transcripts, and protein abundance between Fertile and Sub-fertile heifers from Angus and Holstein genetic backgrounds. Both the independent analysis and multi-omics approach identified potential molecular features capable of discriminating heifers of differing fertility potential.

## METHODS

All analytical procedures are presented in the Additional file 1 and accessible at https://biase-lab.github.io/MultiOmics/

### Ethics Statement

Animal handling for this experiment was approved by the Institutional Animal Care and Use Committee (IACUC) at Virginia Polytechnic Institute and State University.

### Experimental Design

We collected blood samples from purebred Angus heifers (n=12), averaging 14 months in age, at the time of their first artificial insemination (AI) service. Heifers were subjected to a 7-Day Co-Synch + CIDR estrus synchronization protocol prior to breeding. Briefly, heifers were administered an intramuscular (IM) injection of gonadotrophin-releasing hormone (GnRH, 100 μg; Factrel^®^; Zoetis Inc.) on Day 0, followed by the insertion of a controlled internal drug release (CIDR, 1.38 g Progesterone; Eazi-Breed™ CIDR^®^; Zoetis Inc.). On Day 7, the CIDR was removed and an injection of prostaglandin F2 alpha (PGF2α, 25 μg; Lutalyse^®^; Zoetis Inc.) was delivered. Fixed-time AI was performed 54±2 hours following CIDR removal alongside a second injection of GnRH.

Additionally, we collected blood samples from purebred Holstein (n=10) heifers, averaging 12 months in age, at the time of the first AI service. Heifers were enrolled in a 5-Day CIDR-Synch protocol before insemination. Briefly, an IM injection of GnRH was delivered on Day 0 with the insertion of a CIDR device. The CIDR device was removed on Day 5, followed by an IM injection of PGF2α. A second injection of PGF2α was administered 24 hours later. Then, timed AI was performed with a second GnRH injection on Day 8.

Heifers were identified as Fertile (Holstein, n=5; Angus, n=5) or Sub-Fertile (Holstein, n=5; Angus, n=7) based on their pregnancy outcome, following similar criteria used previously [39, 40]. Fertile animals were identified as those who became pregnant and subsequently delivered a calf following the first insemination service. Angus heifers were categorized as Sub-Fertile after failing to achieve pregnancy following two insemination services and exposure to a bull for natural breeding. Holstein heifers were identified as Sub-Fertile after needing four or more artificial inseminations.

### Blood Sample Collection and White Blood Cell Isolation

Fifty ml of blood were drawn from each animal by venipuncture of the jugular vein using 18 mg K2 EDTA vacutainers (Becton, Dickinson, and Company). The tubes were inverted for proper mixing with the anticoagulant and then immediately placed on ice until further processing.

We processed the blood samples following processes described elsewhere [39, 40, 43] within three hours of sampling [43]. Tubes containing whole blood samples were centrifuged for 25 minutes (min) at 4°C and 2,000x*g* to separate the buffy coat. The buffy coat was then aspirated and mixed with 14 ml of red blood cell lysis buffer (1.55 M ammonium chloride, 0.12 M sodium bicarbonate, 1 mM EDTA (Cold Spring Harbor Protocols)). Then, the solution was centrifuged for 10 min at 4°C at 800x*g* and the supernatant was discarded. The remaining pellet was mixed with 200 μl TRIzol™ Reagent (Invitrogen™, Thermo Fisher Scientific, Waltham, MA) in a 2 ml cryotube (Corning Inc., Corning, NY) prior to snap-freezing with liquid nitrogen. Samples were then stored at −80°C until further processing.

### Total RNA and DNA Extraction

The buffy coat samples were thawed at room temperature in a total volume of 525 μl TRIzol™ Reagent. Then, total RNA was extracted from peripheral white blood cells using the Zymo Research Direct-zol™ DNA/RNA Miniprep kit (Zymo Research Corporation, Irvine, CA), according to the manufacturer’s protocol. Next, we assessed the quality of the RNA by quantifying the RNA integrity number (RIN) for each sample using the Agilent RNA 6000 Pico kit (Agilent, Santa Clara, CA) on the Agilent 2100 Bioanalyzer (Agilent, Santa Clara, CA).

### Genotyping and Data Processing

We submitted 400 ng of DNA for each heifer to Neogen (Neogen Corporation, Lincoln, NE) for whole genome sequencing. The samples were genotyped using the Illumina BovineHD Beadchip (Illumina Inc., San Diego, CA) genotyping array (777K). We processed the data for quality control [44] using PLINK [45]. First, we removed SNPs that were preferentially called in one of the groups in the case and control. This was followed by the removal of samples with more than 10% of the genotypes missing, and removal of SNPs with a minor allelic frequency less than 1%, a missing rate greater than 10%, or deviation from the Hardy-Weinberg equilibrium (P<0.00001). Next, we carried out variant pruning. We considered a window size of 50 kilobases with five variants in each window at a correlation threshold of 0.2. All reported SNP coordinates are relative to btau9 assembly converted with the LiftOver tool [46].

### Library Preparation and Sequencing

For sequencing library construction, 900 ng of total RNA was diluted into 25 μl of nuclease-free water, and RNA quantity was confirmed using the Qubit™ RNA High Sensitivity Assay kit (Invitrogen™, Thermo Fisher Scientific, Waltham, MA) on the Qubit™ 4 Fluorometer (Invitrogen™, Thermo Fisher Scientific, Waltham, MA). Libraries were prepared for next-generation sequencing using the Illumina Stranded mRNA Prep kit (Illumina, Inc., San Diego, CA) and the IDT^®^ for Illumina RNA UD indexes (Illumina, Inc., San Diego, CA) according to the manufacturer’s instructions. Sequencing was conducted on the NovaSeq 6000 sequencing system (Illumina, Inc., San Diego, CA) using the NovaSeq 6000 SP Reagent kit v1.5 (Illumina, Inc., San Diego,CA) to produce paired-end reads 150 nucleotides in length. Sequencing was performed by the VANTAGE laboratory at Vanderbilt University Medical Center (Nashville, TN).

### Sequence Alignment and Filtering

We aligned the sequences to the cattle reference genome (Bos_taurus.ARS-UCD1.2.105) in the Ensembl [47] database with hisat2 [48–50] using the -very-sensitive parameter. Then, we used Samtools [51, 52] to filter sequences and remove secondary alignments, duplicates, and unmapped reads. Next, we used biobambam2 [53] to mark and remove duplicates.

### Transcript Quantification and Gene Filtering

The number of fragments that matched to the Ensembl [47] cow gene annotation (Bos_taurus.ARS-UCD1.2.105) was quantified using featureCounts [54], and we preserved genes annotated as protein-coding, pseudogenes, or long non-coding RNA. Genes were then retained for further analysis if counts per million (CPM) and fragments per kilobase per million (FPKM) were >1 in at least five samples.

### Proteomics Data and Processing

One hundred μl of plasma per sample was submitted to the Virginia Tech Mass Spectrometry Incubator (VT-MSI) facility at the Fralin Life Sciences Institute, Virginia Tech, for protein extraction and data collection.

Plasma samples (100 μl) were acidified by the addition of 11.1 μl 12% (v/v) o-phosphoric acid (MilliporeSigma, St. Louis, MO), then proteins were precipitated by the addition of 725 μl LC/MS grade methanol and incubated at −80°C overnight. Precipitated protein was collected by centrifugation and solubilized in S-trap lysis buffer (10% (w/v) SDS in 100 mM triethylammonium bicarbonate ( MilliporeSigma, St. Louis, MO, pH 8.5)). Protein concentration was determined by measuring the absorbance at 280 nm, then 150 μg of protein for each sample was reduced using DTT (4.5 mM) then alkylated with iodoacetamide (10 mM, MilliporeSigma, St. Louis, MO). Unreacted I iodoacetamide was quenched with DTT (10 mM, MilliporeSigma, St. Louis, MO) and samples were acidified using o-phosphoric acid (MilliporeSigma, St. Louis, MO). Protein was again precipitated using methanol and incubated at −80ºC overnight as above. Precipitated protein was loaded onto a micro S-trap and washed with methanol then digested overnight with trypsin. Peptides were recovered and five μg, as determined by measuring the absorbance at 215 nm using a DS-11 FX+ spectrophotometer/fluorometer (DeNovix, Wilmington, DE), of each sample was analyzed twice (duplicates) using ESI-MS/MS Orbitrap Fusion Lumos (Thermo Fisher Scientific (Waltham, MA)).

Samples were first loaded onto a precolumn (Acclaim PepMap 100 (Thermo Scientific, Waltham, MA), 100 μm x 2 cm) after which flow was diverted to an analytical column (50 cm μPAC (PharmaFluidics, Woburn, MA). The UPLC/autosampler utilized was an Easy-nLC 1200 (Thermo Scientific, Waltham, MA). Flow rate was maintained at 150 nl/min and peptides were eluted utilizing a 2 to 45% gradient of solvent B in solvent A over 88 minutes. Spray voltage on the μPAC compatible Easy-Spray emitter (PharmaFluidics, Woburn, MA) was 1300 volts, the ion transfer tube was maintained at 275°C, the RF lens was set to 30% and the default charge state was set to 3.

MS data for the m/z range of 400-1500 was collected using the orbitrap at 120000 resolution in positive profile mode with an AGC target of 4.0e5 and a maximum injection time of 50 ms. Peaks were filtered for MS/MS analysis based on having isotopic peak distribution expected of a peptide with an intensity above 2.0e4 and a charge state of 2-5. Peaks were excluded dynamically for 15 seconds after 1 scan with the MS/MS set to be collected at 45% of a chromatographic peak width with an expected peak width (FWHM) of 15 seconds. MS/MS data starting at m/z of 150 was collected using the orbitrap at 15000 resolution in positive centroid mode with an AGC target of 1.0e5 and a maximum injection time of 200 ms. Activation type was HCD stepped from 27 to 33.

Data were analyzed utilizing Proteome Discoverer 2.5 (Thermo Scientific, Waltham, MA) combining a Sequest HT and Mascot 2.7 (Matrix Science, Boston, MA) search into one result summary for each sample. Both searches utilized the UniProt reference *Bos taurus* proteome database and a common protein contaminant database provided with the Proteome Discoverer (PD) software package [55]. Each search assumed trypsin-specific peptides with the possibility of 2 missed cleavages, a precursor mass tolerance of 10 ppm and a fragment mass tolerance of 0.1 Da. Sequest HT searches also included the PD software precursor detector node to identify MS/MS spectra containing peaks from more than one precursor. Sequest HT searches included a fixed modification of carbamidomethyl at Cys and the variable modifications of oxidation at Met and loss of Met at the N-terminus of a protein (required for using the INFERYS rescoring node). Peptide matches identified by Sequest HT were subjected to INFERYS rescoring to further optimize the number of peptides identified with high confidence.

Mascot searches included the following dynamic modifications in addition to the fixed modification of Cys alkylated by iodoacetamide (carbamidomethylated): oxidation of Met, acetylation of the protein N-terminus, cyclization of a peptide N-terminal Gln to pyro-Glu, and deamidation of Asn/Gln residues.

Protein identifications were reported at a 1% false discovery rate (high confidence) or at 5% false discovery rate (medium confidence) based on searches of decoy databases utilizing the same parameters as above. The software matched peptide peaks across all runs, and protein quantities are the sum of all peptide intensities associated with the protein.

### Statistical Analyses

#### SNP Association Analysis

After filtering, 575,053 genotypes from 22 animals were used for association analysis conducted in PLINK [45] using Fisher’s exact test. Locus association was inferred at alpha = 1×10^-5^, as reported by The Wellcome Trust Case Control Consortium [56] for case-control studies and previous GWAS analyses of reproductive traits in cows or heifers [4, 25, 32].

#### Differential Transcript Abundance

We compared transcript abundance between samples from each breed and each fertility group. The R packages ‘edgeR’ [57, 58], with the quasi-likelihood test, and ‘DEseq2’ [59], using the Wald’s and likelihood test, were utilized to conduct the analyses. We adjusted the raw *P* values for multiple hypothesis testing by calculating the empirical false discovery rate (eFDR, [60]) with 10,000 permutations. Differences in transcript abundance were deemed statistically significant when eFDR <0.002 in the results obtained from the three tests.

#### Differential Protein Abundance

To identify differential protein abundance that is robust to the algorithm utilized, we analyzed the protein data using two different algorithms. First, we transformed the protein data using natural logarithm (Log_*e*_(x)). We analyzed the transformed data using a generalized mixed model [61] using the R package ‘lme4’, which included the fertility group (Fertile or Sub-Fertile), breed (Angus or Holstein), and the random effect of the subject. Random effect was included in this analysis as samples were assayed twice to provide a more robust estimate of differential protein abundance. Then, we used the function “emmeans”, which tests the significance of the difference (H_0_:μ_1_=μ_2_, H_1_:μ_1_≠μ_2_) with the Student’s t test [62], to calculate the estimated differences in protein abundance between fertility groups within each breed. We also analyzed the log-transformed data using the R package ‘limma’ [63]. We accounted for the same independent variables mentioned above (fertility group and breed), in addition to accounting for the correlation between the duplicated data for each individual with the function ‘duplicateCorrelation’. We tested for a differential abundance of the identified proteins using the empirical Bayes Statistics implemented in the function ‘eBayes’ [64, 65]. In both analyses, we adjusted the nominal P values using FDR [66]. Significance was assumed if FDR< 0.05 in both approaches.

#### Multi-omics Factor Analysis

We analyzed the multimodal multi-omics datasets (genome, transcriptome and proteome) interactively using Multi-omics Factor Analysis approach [67, 68]. We subset the genotypes, transcriptome, and proteome data to reduce the global profiling. We retained SNPs with a P value < 0.001 for the Fisher’s test, genes with a P value < 0.01 for all three statistical tests employed, and proteins with a P value < 0.05 in both statistical tests used. We conducted the analysis using the R package “MOFA2” [67, 68], accounting for the breed as a group.

## RESULTS

### Overview of the data produced

We selected 22 *Bos taurus* heifers of Angus (n=12) and Holstein (n=10) breeds based on their fertility fitness (Figure 1A). We isolated total RNA from circulating white blood cells, averaging 16.3 μg±4.0, and quality, measured by the RIN, averaging 9.4±0.4. The extraction of genomic DNA yielded 1.1 μg±0.4. We produced RNA-sequencing data (Figure 1B) and quantified the transcript abundance of 12,445 genes (12,105 protein-coding genes, 228 long non-coding RNAs, and 112 pseudogenes). We also analyzed 575,053 nucleotide positions across the bovine genome (Figure 1B). Lastly, we produced untargeted proteomics data from plasma that resulted in the relative quantification of 213 proteins. As expected, the genotypic and proteomic data clustered the heifers of different genetic background separately. By contrast, there was no clustering of the samples based on the transcriptome data of the peripheral white blood cells (Figure 1 C,D,E).

**Figure 1.**
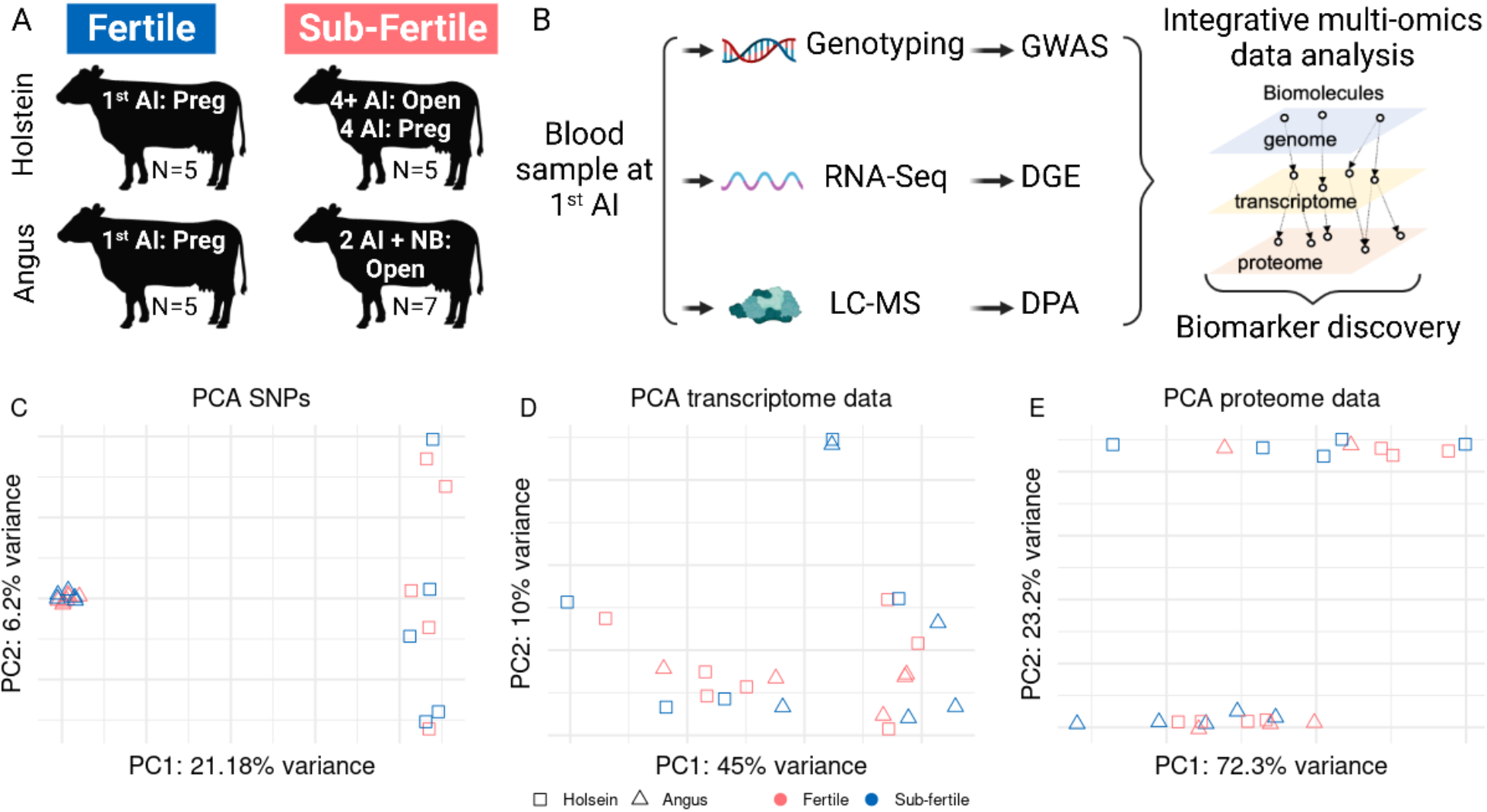
Overview of data produced. (A) Breeds and classification used in this study, including sample size. (B) Schematics of the data produced, and analysis undertaken. Principal component analysis of the genome-wide single nucleotide polymorphisms (C), transcriptome (D) and (E) proteome data. GWAS: genome-wide association analysis, DGE: differential gene expression, DPA: differential protein abundance

### GWAS identifies SNPs associated with fertility in Angus and Holsteins heifers

Our analysis identified two SNPs significantly associated with heifer fertility (rs110918927, chr12: 85648422, P = 6.7×10^-7^; and rs109366560, chr11:37666527, P = 2.6×10^-5^, Figure 2A, Additional file 2). For the SNP rs110918927, all heifers that delivered a calf after one artificial insemination presented genotype AA (f_(A)_=1, f_(G)_=0), whereas 11 out of 12 heifers classified as sub-fertile presented at least one copy of the allele G (f_(A)_=0.29, f_(G)_=0.71). For the SNP rs109366560, none of the heifers classified as sub-fertile were homozygous for the allele G (f_(G)_=0.12, f_(A)_=0.88), and five out of the nine fertile heifers genotyped were homozygous GG (f_(G)_=0.78, f_(A)_=0.22).

**Figure 2.**
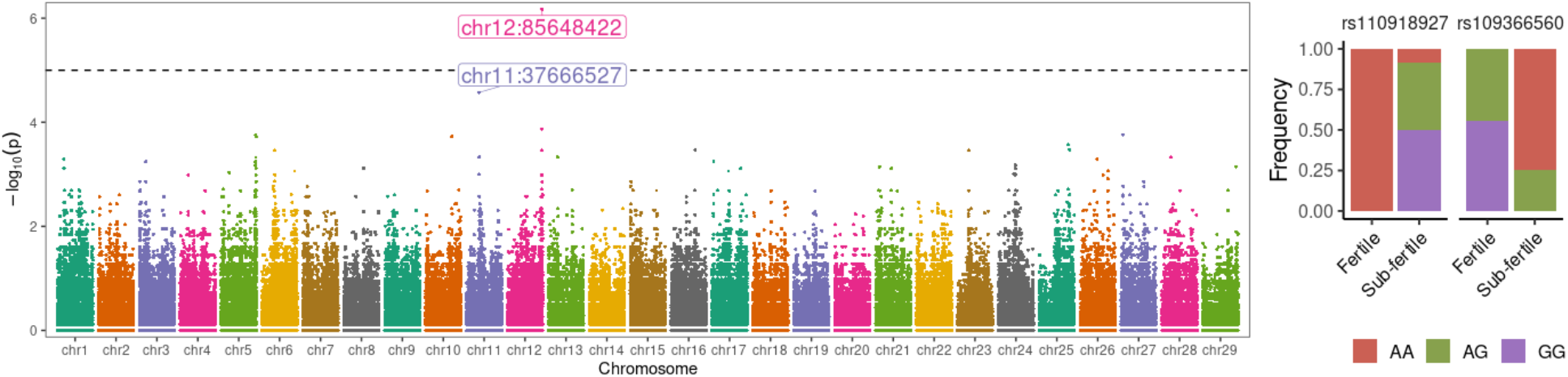
Genome-wide association analysis of fertility in beef and dairy heifers. (A) Manhattan plot with the distribution of SNPs across their genome and their P values from Fisher’s exact association test. (B) Genetic frequencies of the two SNPs that are putatively linked to fertility in heifers.

### Transcriptome analysis identifies differential transcript abundance between Fertile and Sub-Fertile heifers

Next, we sought to determine if there were differences in transcript abundance from circulating white blood cells between the Fertile and Sub-Fertile heifer groups, accounting for their genetic background. We identified two genes whose transcript abundance differed (eFDR ≤ 0.002) between the two groups (Fertile and Sub-Fertile), namely Adipocyte Plasma Membrane Associated Protein (*APMAP*, 1.16 greater abundance in the Fertile group) and Dynein Axonemal Intermediate Chain 7 (*DNAI7*, 1.23 greater abundance in the Sub-Fertile group) (Additional file 3).

### Proteomic analysis identifies differential protein abundance between Fertile and Sub-Fertile heifers

We also tested if there were differential abundance in proteins present in the plasma of heifers classified based on their fertility groups in both genetic backgrounds. The protein Alpha-ketoglutarate-dependent dioxygenase FTO was more abundant in the plasma collected from Fertile heifers relative to their Sub-Fertile counterparts (FDR < 0.05, Figure 4A). Two other proteins did not reach the significance threshold in the model accounting for all samples (Apolipoprotein C-II, APOC2 (FDR_glmm_ = 0.06) and Lymphocyte cytosolic protein 1, LCP1 (FDR_glmm_ = 0.06)), but the within-breed estimates showed a lower abundance of these proteins in Fertile Holstein heifers (P < 0.05, Figure 4B, C) (Additional file 4).

**Figure 3.**
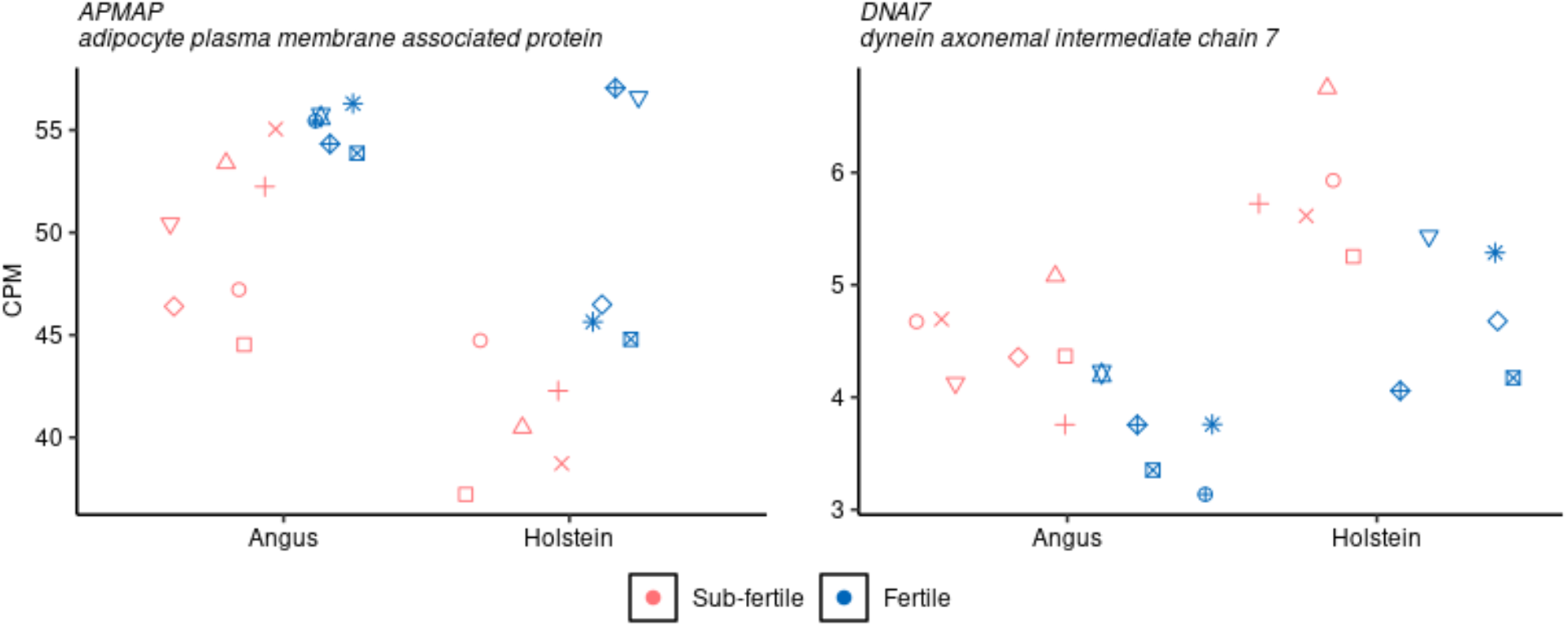
Differential transcript abundance associated with fertility. In each plot, within each breed, shapes indicate same animal.

**Figure 4.**
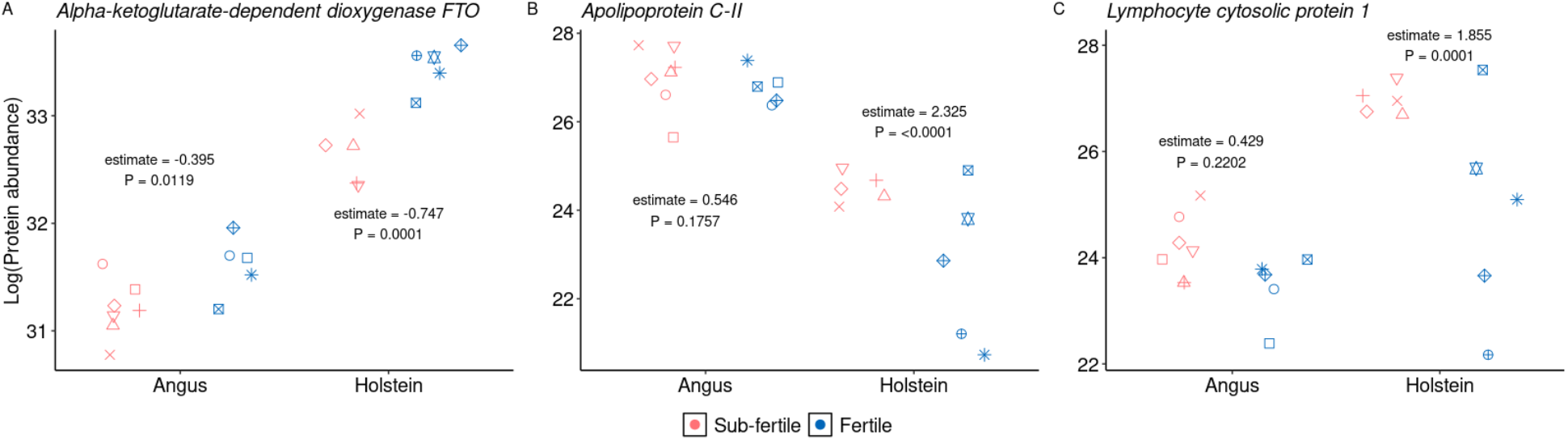
Differential protein abundance associated with fertility. In each plot, within each breed, shapes indicate same animal. Within each breed, estimates are for the contrast Sub-Fertile – Fertile heifers.

### Integrative multi-omics analysis identifies molecular features

When each data were evaluated independently, the quantification of 22 and 23 gene and protein relative abundances and the genotypic information of 59 SNPs explained 44.1%, 16.6%, and 70.1% of the variance associated with fertility classification, respectively. Overall, there were four factors identified in the analysis with the potential to distinguish the samples based on their fertility status, out of which three were most representative with Factors one, two, and three being mostly dominated by genotype, transcript, and protein data, respectively (Figure 5A). Factors one, two, and three separated most of the samples based on their fertility classification except two, three, and five samples, respectively (Figure 5B).

**Figure 5.**
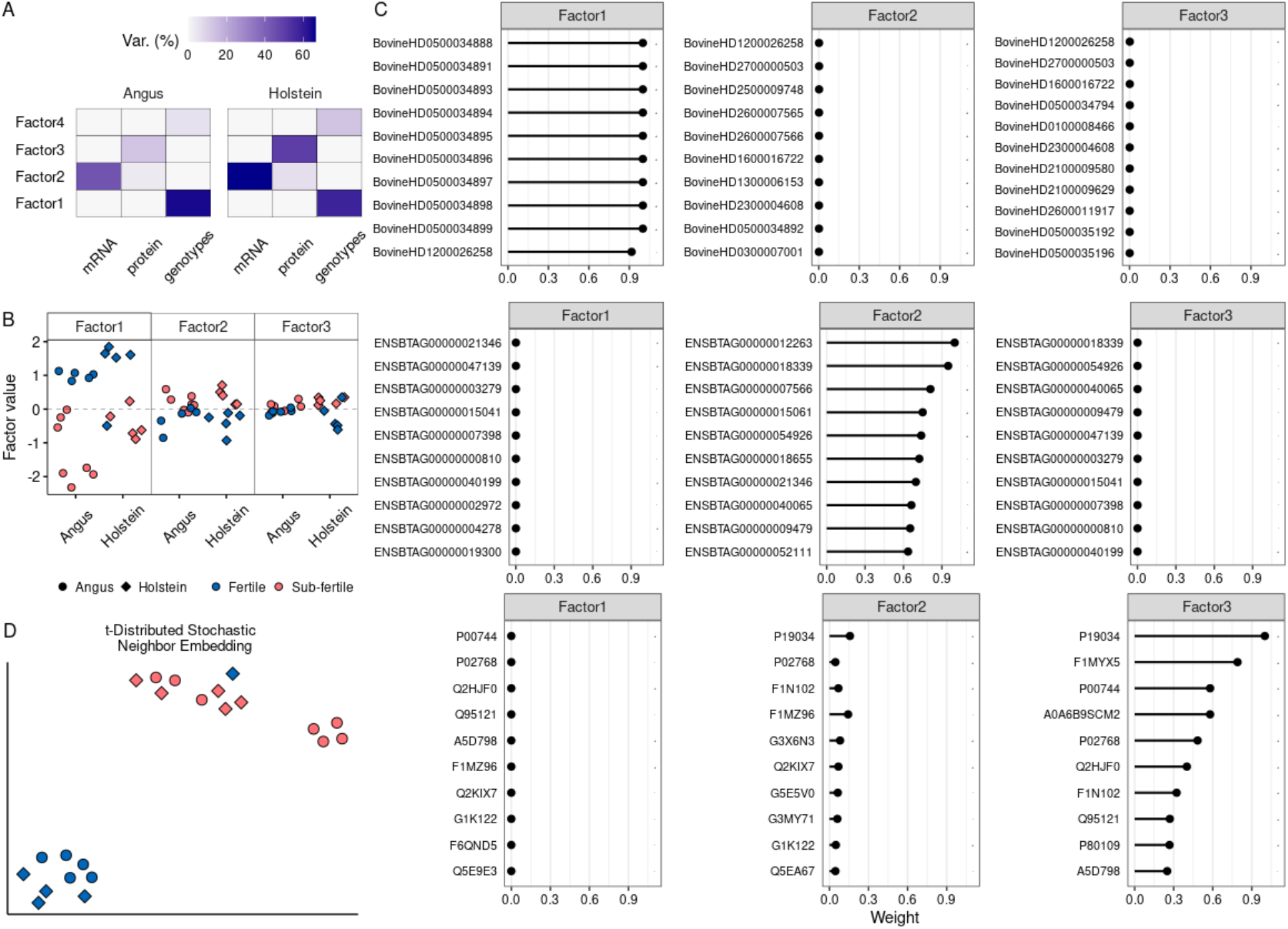
Multi-omics analysis of heifer fertility. (A) Percentage of variance explained by each factor within each data modality. (B) Relative separation of samples based on their phenotype for each factor plotted. (C) The relative weight of importance of the top ten features (SNP identifiers are from the array annotation, gene identifiers are from Ensembl, protein identifiers are from UniProt) in each data modality and factor plotted.

Notably, the top nine SNPs that explained most of the variance related to Factor one encompass a window in chromosome 5 spanning nucleotides 118332762 and 118345383. The tenth SNP was the top significant polymorphism identified on chromosome 12 nucleotide 85648422 according to our Fisher’s exact test contrasting heifers of different fertility potential (Figure 5C). Among the genes whose transcript abundance explained the variance related to Factor two, we identified the following annotated genes: ArfGAP with SH3 domain, ankyrin repeat and PH domain 3 (*ASAP3*), ATP synthase membrane subunit c locus 1 (*ATP5MC1*), Centrosomal protein 170 (*CEP170*), Myeloid derived growth factor (*MYDGF*), Coiled-coil domain containing 34 (*CCDC34*), RAD51 associated protein 1 (*RAD51AP1*), Ubiquinol-cytochrome c reductase complex III subunit VII (*UQCRQ*) (Figure 5C). Among the proteins whose abundance explained the variance related to Factor three, the following were annotated to known genes: Apolipoprotein C-II (*APOC2*), Lymphocyte cytosolic protein 1 (*LCP1*), Vitamin K-dependent protein Z (*PROZ*), Albumin (*ALB*), Serotransferrin-like (LOC525947), Complement component C8 beta chain (*C8B*), Pigment epithelium-derived factor (*SERPINF1*), Phosphatidylinositol-glycan-specific phospholipase D (*GPLD1*), Alpha-ketoglutarate-dependent dioxygenase FTO (*FTO*) (Figure 5C). Collectively, data from genotypes, transcriptome, and proteome clustered 21 out of 22 heifers correctly based on their fertility status, with only one Fertile heifer clustering with the group of Sub-Fertile heifers.

## Discussion

Reproduction is a multidimensional biological function in mammals that can be partitioned into multiple components or traits [69], and as a consequence, infertility is a complex phenotype with multifactorial origins. Our work addressed two critical questions: 1-whether multiple layers of molecular information, present in the circulatory system, would be associated with female fertility fitness; and 2-whether the integrative analysis of multiple layers of molecular information would be a better predictor of heifers with different fertility potentials. Our analysis identified molecular signatures in the genome, transcriptome, and proteome associated with heifer infertility. More importantly, the integration of multiple levels of biological information produces a signature capable of separating most fertile from the sub-fertile heifers.

The top polymorphism (rs110918927) significantly associated with fertility located on chr12:85648422 sits in an intergenic region of the genome with the closest gene located >73 kilobases downstream relative to the SNP. The second significant polymorphism (rs109366560) is located on an intron (chr11:37666527) of the gene Echinoderm microtubule associated protein like 6 (*EML6*). Neither SNP is located in a region previously associated with female reproductive traits [70]. These SNPs have also not been previously reported to be associated with fertility traits in previous investigations that focused on sire-centric models [4, 33–37], nor on studies that focused on genotyped heifers only [32, 38].

It is notable that the gene *EML6* produces a protein functionally associated with the spindle microtubules in oocytes [71]. Knockdown of this protein in mice oocytes at the germinal vesicle stage impairs spindle morphology and increases aneuploidy in oocytes that progress to the metaphase II stage [71]. The gene *EML6* also produces transcripts in bovine oocytes [72], and the significant SNP in this gene is a strong indication of a functional connection to reduced oocyte developmental competence in the Sub-Fertile group of heifers

Genes differentially expressed in the peripheral white blood cells have been associated with fertility in heifers [39–41]. The protein APMAP exhibits arylesterase activity, which is known to protect lipoproteins from oxidation [73]. The protein APMAP regulates adipose composition and metabolic health and the disruption of the *APMAP* gene in mice leads to an increase in visceral adipose tissue expansion [74]. The protein APMAP was less abundant in the omental tissue of women diagnosed with polycystic ovary syndrome [75]. Therefore, lower expression of *APMAP* in the peripheral white blood cells of Sub-Fertile heifers is possibly connected with a metabolic, hormonal or inflammatory disorder that disrupts fertility in heifers.

The Protein DNAI7 composes the axonemal dynein complex and participates in beta-tubulin binding activity and microtubule binding activity, and thus contribute to ciliary beating [76]. Variants that impair the function of DNAI7 are associated with Primary Ciliary Dyskenesia, with one potential consequence being the abnormal function of cilia and possible impaired transport of the cleaving embryos into the uterus [77]. DNAI7 may also function as a cell cycle regulator, and dysregulated transcript abundance of DNAI7 was associated with nasopharyngeal neoplasm in mice [78] and lung adenocarcinoma in humans [79]. Because Sub-Fertile heifers have greater abundance of *DNAI7* transcripts in their circulating white blood cells, it is possible that dysregulation in the cell cycle has a biological link with subfertility. Further research is required, however, to evaluate whether a dysregulation in the cell cycle linked to upregulation of *DNAI7* is connected with increased inflammation [74] associated with less transcripts from *APMAP*.

The protein Alpha-ketoglutarate-dependent dioxygenase FTO has an oxidative demethylation activity of abundant N6-methyladenosine (m^6^A) residues in RNA [80]. The protein FTO preferentially demethylates N6,2’-O-dimethyladenosine (m^6^A_m_) rather than m^6^A and contributes to a reduced stability of m^6^A_m_ mRNAs [81]. On a systemic level, genomic variants in the *FTO* were associated with symptoms of metabolic disorders [82], although the effects observed in humans, such elevated body mass index [83, 84], and mice [85] may be contradictory. Also worth noting, a variant on the *FTO* gene was associated with polycystic ovary syndrome [86]. Interestingly, in mice, the *FTO* gene is downregulated due to deficiency in essential amino-acids [87], and deficiency in the FTO protein causes postnatal growth retardation and a significant reduction in adipose tissue and lean body mass [88]. Our observation of the FTO abundance in heifers of different fertility potential is an indication that Sub-Fertile heifers could be experiencing metabolic imbalance, contributing to their lower fertility.

We must highlight the results observed for Apolipoprotein C-II and Lymphocyte cytosolic protein 1. Both proteins had a greater abundance in Sub-Fertile Holstein heifers. The major function of the APOC2 protein is to support the activation of lipoprotein lipase, which in turn hydrolyzes triglycerides making fatty acids available to cells [89]. In mice, the ApoC2 gene is upregulated by high lipid load [90]. In humans, APOC2 is highly abundant in the serum of patients with cancer [91–94], and the current hypothesis is that APOC2 provides the lipid metabolism, and thus the energy needed for tumorigenesis and tumor progression [94, 95]. The protein LCP1 is an actin-binding protein [96] that contributes to actin interaction and assembly in cells [97]. Despite the seemingly structural function, the protein LCP1 is associated with the differentiation, proliferation, and migration of immune cells [98–102]. The higher expression of APOC2 and LCP1 collectively add to the indication of a possible metabolic or inflammatory disorder linked with heifer subfertility in Holsteins.

The next step was to interrogate the data we produced in a comprehensive manner. Interestingly, the largest source of variability was observed in the genomic data. Nine of the top ten SNPs that were assigned to Factor one were located in an intron of the TAFA chemokine-like family member 5 (*TAFA5*) gene. These SNPs are within a quantitative trait loci for milk yield [103], a trait negatively correlated with reproduction traits [104], however, no relationship between genetic variants in this gene and female fertility has been reported previously. None of the top ten genes with transcript abundance relevant for the modeling of the variance were identified as differentially expressed when analyzed independently. This result is not surprising because the identification of significant features using standard statistical approaches for association analysis is not necessarily the best approach for identifying predictive genes associated with complex traits [105, 106]. It was surprising that three out of nine annotated proteins, which composed the top ten proteins that explained most of the variance in factor three, were also identified in our analyses using general linear mixed models. The most interesting result, however, was that all three data modalities were able to separate 21 out of 22 heifers correctly based on their fertility potential. Our results support the biological complexity involving female infertility, but also shed light on molecular differences with important biological insights for our comprehensive understanding of female infertility.

## Conclusion

Our interrogation of multiple levels of biological information (genome, transcriptome, and proteome) at a systemic level in heifers highlight the complexity of female fertility. While the genomic data point to a disruption of oocyte developmental competence, the transcriptome and proteomic data point to a metabolic dysregulation contributing to subfertility. Lastly, our results highlight differences in the molecular profile of heifers associated with fertility that transcend the constraints of breed-specific genetic background.

## List of abbreviations

μg: microgram
AI: artificial insemination
*ALB*: Albumin
*APMAP*: Adipocyte Plasma Membrane Associated Protein
APOC2: Apolipoprotein C-II
*ASAP3*: ArfGAP with SH3 domain, ankyrin repeat and PH domain 3
*ATP5MC1*: ATP synthase membrane subunit c locus 1
*C8B*: Complement component C8 beta chain
*CCDC34*: Coiled-coil domain containing 34
*CEP170*: Centrosomal protein 170
CIDR: controlled internal drug release
CPM: counts per million
*DNAI7*: Dynein Axonemal Intermediate Chain 7
DTT: dithiothreitol
DGE: differential gene expression
DPA: differential protein abundance
eFDR: empirical false discovery rate
*EML6*: Echinoderm microtubule associated protein like 6
FDR: false discovery rate
FPKM: fragments per kilobase per million
*FTO*: Alpha-ketoglutarate-dependent dioxygenase FTO
GnRH: gonadotrophin-releasing hormone
*GPLD1*: Phosphatidylinositol-glycan-specific phospholipase D
GWAS: genome-wide association study
IAA: indole-3-acetic acid
IACUC: Institutional Animal Care and Use Committee
IM: intramuscular
K2 EDTA: dipotassium ethylenediaminetetraacetic acid
LCP1: Lymphocyte cytosolic protein 1
LOC525947: Serotransferrin-like
M: molar
m^6^A: N6-methyladenosine
m^6^A_m_: N6,2’-O-dimethyladenosine
mg: milligram
Min: minutes
ml: milliliter
mRNA: messenger RNA
ms: millisecond
*MYDGF*: Myeloid derived growth factor
ng: nanogram
nm: nanometer
nl: nanoliter
P: probability
PGF2α: prostaglandin F2 alpha
*PROZ*: Vitamin K-dependent protein Z
*RAD51AP1*: RAD51 associated protein 1
RIN: RNA integrity number
*SERPINF1*: Pigment epithelium-derived factor
SNP: single nucleotide polymorphism
TAFA5: TAFA chemokine-like family member 5
*UQCRQ*: Ubiquinol-cytochrome c reductase complex III subunit VII
VT-MSI: Virginia Tech Mass Spectrometry Incubator
*Xg*: relative centrifugal force
μl: microliter

## Declarations

### Consent for publication

Not applicable

### Availability of data and material

The transcriptome and proteome data generated and analyzed during the current study are available in the Gene Expression Omnibus and ProteomeXchange repositories under the following identifiers: GSE220220 and PXD038756, respectively. The genotypic data are available from the corresponding author upon reasonable request.

### Authors’ contributions

FB designed, supervised, and acquired funding. FB collected and processed the samples for data collection. MM contributed to funding acquisition, analysis, and interpretation of the data. FB and MM worked on the draft. All authors have read and approved the submitted version.

## Acknowledgments

We express our greatest appreciation for Chad Joines, Director of Beef Cattle Operations, Shane Brannock, Director of Dairy Complex, and their respective crews for supporting our sampling. We also thank Jada Nix from our group for the critical reading of our manuscript.

## Supplementary Material

File name: Additional file 1

File format: html

Title of data: Supplementary code to: A multi-omics analysis identifies molecular features associated with heifer fertility in a case-control design including Angus and Holstein cattle

Description of data: The code utilized for analytical procedures to obtain the results and graphs.

File name: Additional file 2

File format: .xlsx

Title of data: Genome-wide association analysis of heifer fertility

Description of data: Results of Fisher’s exact test of GWAS analysis of heifer fertility.

File name: Additional file 3

File format: .xlsx

Title of data: Results of differential transcript abundance associated with heifer fertility

Description of data: Results of differential transcript abundance associated with heifer fertility

File name: Additional file 4

File format: .xlsx

Title of data: Results of differential protein abundance associated with heifer fertility

Description of data: Results of differential protein abundance associated with heifer fertility

