## Additional file 1 for "A multi-omics analysis identifies molecular features associated with heifer fertility in a case-control design including Angus and Holstein cattle"

#### 2022-12-13


### Overview

Code produced by Fernando Biase, and Mackenzie Marrella. We created
this file to permit reproducibility of the findings described in the
paper. Please direct questions to Fernando Biase:
***fbiase*** at ***vt.edu***

The raw transcriptome data is deposited on GEO repository under the
following access GSE220220.
The normalized protein data obtained from the facility is deposited in
the ProteomeXchange repository under the following access PXD038756.

Please contact Fernando Biase: ***fbiase*** at
***vt.edu*** if you’d like access to other files
used in this code. Updated information may be obtained at www.biaselaboatory.com

**ABSTRACT**: **Background**: Infertility
or subfertility is a critical barrier to sustainable cattle production,
including in heifers. The development of heifers that do not produce a
calf within an optimum window of time is a critical factor for the
profitability and sustainability of the cattle industry. The early
identification of heifers with optimum fertility using molecular
phenotyping is a promising approach to improving sustainability in beef
and dairy cattle production.  
 **Results**: Using a
high-density SNP chip, we collected genotypic data from 575,053 SNPs. We
also produced quantitative transcriptome data for 12,445 genes (12,105
protein-coding genes, 228 long non-coding RNAs, and 112 pseudogenes) and
proteome data for 213 proteins. We identified two SNPs significantly
associated with heifer fertility (rs110918927, chr12: 85648422, P =
6.7x10-7; and rs109366560, chr11:37666527, P = 2.6x10-5). We identified
two genes with differential transcript abundance (eFDR ≤ 0.002) between
the two groups (Fertile and Sub-Fertile): Adipocyte Plasma Membrane
Associated Protein (APMAP, 1.16 greater abundance in the Fertile group)
and Dynein Axonemal Intermediate Chain 7 (DNAI7, 1.23 greater abundance
in the Sub-Fertile group). Our analysis revealed that the protein
Alpha-ketoglutarate-dependent dioxygenase FTO was more abundant in the
plasma collected from Fertile heifers relative to their Sub-Fertile
counterparts (FDR < 0.05). Interestingly, two proteins did not reach
the significance threshold in the model accounting for all samples
(Apolipoprotein C-II, APOC2 (FDRglmm = 0.06) and Lymphocyte cytosolic
protein 1, LCP1 (FDRglmm = 0.06)), but both proteins were less abundant
in the plasma of Fertile Holstein heifers (P < 0.05). Lastly, an
integrative analysis of the three datasets identified a series of
features (SNPs, gene transcripts, and proteins) that can be useful for
the discrimination of heifers based on their fertility. When all
features were utilized together, 21 out of 22 heifers were classified
correctly based on their fertility category.  
**Conclusions**: Our multi-omics analyses confirm the
complex nature of female fertility. Very importantly, our results also
highlight differences in the molecular profile of heifers associated
with fertility that transcend the constraints of breed-specific genetic
background.

---

```
library('dplyr', quietly = TRUE,lib.loc="/usr/lib/R/site-library")
library('ggplot2', quietly = TRUE,lib.loc="/usr/lib/R/site-library")
library('edgeR',quietly = TRUE, lib.loc="/usr/lib/R/site-library")
library("cowplot", quietly = TRUE,lib.loc="/usr/lib/R/site-library")
library("GGally",quietly = TRUE,lib.loc="/usr/lib/R/site-library")
library("DESeq2",quietly = TRUE,lib.loc="/usr/lib/R/site-library")
library('readxl',quietly = TRUE,lib.loc="/usr/lib/R/site-library")
library("VennDiagram", quietly = TRUE, lib.loc="/usr/lib/R/site-library")
library("ComplexHeatmap",quietly = TRUE, lib.loc="/usr/lib/R/site-library")
library("flashClust", quietly = TRUE, lib.loc="/usr/lib/R/site-library")
library('plotly' , quietly = TRUE, lib.loc="/usr/lib/R/site-library")
library('tidyverse' ,quietly = TRUE , lib.loc="/usr/lib/R/site-library")
library('htmlwidgets' , quietly = TRUE, lib.loc="/usr/lib/R/site-library")
library('reshape2' ,quietly = TRUE, lib.loc="/usr/lib/R/site-library")
library("ggpubr" ,quietly = TRUE, lib.loc="/usr/lib/R/site-library")
library("car" ,quietly = TRUE, lib.loc="/usr/lib/R/site-library") 
library("goseq" ,quietly = TRUE, lib.loc="/usr/lib/R/site-library")
library("stringr",quietly = TRUE , lib.loc="/usr/lib/R/site-library")
library("data.table",quietly = TRUE , lib.loc="/usr/lib/R/site-library")
library("tidyr",quietly = TRUE , lib.loc="/usr/lib/R/site-library")
library("dplyr",quietly = TRUE , lib.loc="/usr/lib/R/site-library")
library("ggsignif",quietly = TRUE , lib.loc="/usr/lib/R/site-library")
library("kableExtra",quietly = TRUE , lib.loc="/usr/lib/R/site-library")
library("grid",quietly = TRUE , lib.loc="/usr/lib/R/site-library")
library("gridExtra",quietly = TRUE , lib.loc="/usr/lib/R/site-library")
library("bigmemory",quietly = TRUE , lib.loc="/usr/lib/R/site-library")
library("doParallel",quietly = TRUE , lib.loc="/usr/lib/R/site-library")
library("ggmanh",quietly = TRUE , lib.loc="/usr/lib/R/site-library")
library("lme4",quietly = TRUE , lib.loc="/usr/lib/R/site-library")
library("emmeans",quietly = TRUE , lib.loc="/usr/lib/R/site-library")
library("limma",quietly = TRUE , lib.loc="/usr/lib/R/site-library") 
library("MultiAssayExperiment",quietly = TRUE , lib.loc="/usr/lib/R/site-library")
library('MOFA2', lib.loc="/usr/lib/R/site-library")
```

### Procedures

Use the tabs below to access each section of our code.

#### Get resources for reproducibility

##### Obtain Ensembl Annotation

```
library('biomaRt', lib.loc="/usr/lib/R/site-library")
cow<-useMart("ensembl", dataset = "btaurus_gene_ensembl", host="www.ensembl.org") 
annotation.ensembl.symbol<-getBM(attributes = c('ensembl_gene_id','external_gene_name','description','hgnc_symbol','gene_biotype','transcript_length','chromosome_name','start_position','end_position','strand'),  values = "*", mart = cow)

cow<-useMart("ensembl", dataset = "btaurus_gene_ensembl", host="www.ensembl.org") 
annotation.ensembl.transcript<-getBM(attributes = c('ensembl_gene_id','gene_biotype','ensembl_transcript_id','transcript_length'),  values = "*", mart = cow)
annotation.ensembl.transcript<-annotation.ensembl.transcript[annotation.ensembl.transcript$gene_biotype =="protein_coding",]
annotation.ensembl.transcript<-annotation.ensembl.transcript[order(annotation.ensembl.transcript$ensembl_gene_id, -annotation.ensembl.transcript$transcript_length),]
annotation.ensembl.transcript<-annotation.ensembl.transcript[!duplicated(annotation.ensembl.transcript$ensembl_gene_id),]
annotation.ensembl.transcript<-annotation.ensembl.transcript[annotation.ensembl.transcript$transcript_length > 400,]

annotation.ensembl.symbol<-annotation.ensembl.symbol[order(annotation.ensembl.symbol$ensembl_gene_id, -annotation.ensembl.symbol$transcript_length),]
annotation.ensembl.symbol<-annotation.ensembl.symbol[!duplicated(annotation.ensembl.symbol$ensembl_gene_id),]
gene.length<-annotation.ensembl.symbol[,c( "ensembl_gene_id", "transcript_length" )]
annotation.GO.biomart<-getBM(attributes = c('ensembl_gene_id', 'external_gene_name','go_id','name_1006','namespace_1003'),  values = "*", mart = cow)

#write.table(annotation.ensembl.symbol,file="/mnt/storage/lab_folder/shared_R_codes/chace/angus_holstein_AI/resources/2022_05_25_annotation.ensembl.symbol.txt", sep = "\t",append = FALSE, quote = FALSE)
#system('bzip2 --best /mnt/storage/lab_folder/shared_R_codes/chace/angus_holstein_AI/resources/2022_05_25_annotation.ensembl.symbol.txt')

#write.table(gene.length,file="/mnt/storage/lab_folder/shared_R_codes/chace/angus_holstein_AI/resources/2022_05_25_gene.length.txt", sep = "\t",append = FALSE, quote = FALSE)
#system('bzip2 --best /mnt/storage/lab_folder/shared_R_codes/chace/angus_holstein_AI/resources/2022_05_25_gene.length.txt')

#write.table(annotation.GO.biomart,file="/mnt/storage/lab_folder/shared_R_codes/chace/angus_holstein_AI/resources/2022_05_25_annotation.GO.biomart.txt", sep = "\t",append = FALSE, quote = FALSE)
#system('bzip2 --best /mnt/storage/lab_folder/shared_R_codes/chace/angus_holstein_AI/resources/2022_05_25_annotation.GO.biomart.txt')

#write.table(annotation.ensembl.transcript,file="/mnt/storage/lab_folder/shared_R_codes/chace/angus_holstein_AI/resources/2022_05_25_annotation.ensembl.transcript.txt", sep = "\t",append = FALSE, quote = FALSE)
#system('bzip2 --best /mnt/storage/lab_folder/shared_R_codes/chace/angus_holstein_AI/resources/2022_05_25_annotation.ensembl.transcript.txt')
```

##### Import Annotation

```
annotation.ensembl.symbol<-read.delim("/mnt/storage/lab_folder/shared_R_codes/chace/angus_holstein_AI/resources/2022_05_25_annotation.ensembl.symbol.txt.bz2", header=TRUE, sep= "\t",row.names=1, stringsAsFactors = FALSE)
gene.length<-read.delim("/mnt/storage/lab_folder/shared_R_codes/chace/angus_holstein_AI/resources/2022_05_25_gene.length.txt.bz2", header=TRUE, sep= "\t",row.names=1, stringsAsFactors = FALSE)
annotation.GO.biomart<-read.delim("/mnt/storage/lab_folder/shared_R_codes/chace/angus_holstein_AI/resources/2022_05_25_annotation.GO.biomart.txt.bz2", header=TRUE, sep= "\t",row.names=1, stringsAsFactors = FALSE)
annotation.ensembl.transcript<-read.delim("/mnt/storage/lab_folder/shared_R_codes/chace/angus_holstein_AI/resources/2022_05_25_annotation.ensembl.transcript.txt.bz2", header=TRUE, sep= "\t",row.names=1, stringsAsFactors = FALSE)
```

#### GWAS analysis

##### Import raw data and prepare files for Plink

```
#set up phenotype data for analysis
snp_map_genotypes <- read.table("/mnt/storage/lab_folder/heifer_infertility/AI_angus_holstein/assoc_analysis/SNP_Map.txt", header = TRUE, sep = "\t")
ah_genotypes<- data.table::fread("/mnt/storage/lab_folder/heifer_infertility/AI_angus_holstein/assoc_analysis/Virginia_Tech_Univ_Biase_BOV770V01_20220825_FinalReport.txt", header = TRUE, sep = "\t", skip = 9)

ah_genotypes$Sample.ID <- gsub(" ","_",ah_genotypes$Sample.ID)
ah_genotypes <- ah_genotypes[,c(2,2,1,3,4)]
ah_genotypes$Allele1...Forward <- gsub("-","0",ah_genotypes$Allele1...Forward)
ah_genotypes$Allele2...Forward <- gsub("-","0",ah_genotypes$Allele2...Forward)
ah_genotypes<- ah_genotypes[!ah_genotypes$Sample.ID=="Sample_23" & !ah_genotypes$Sample.ID=="Sample_24",]


snp_map_genotypes <- snp_map_genotypes[,c(3,2,4)]


sample_id <- c("Sample_1","Sample_2","Sample_3","Sample_4","Sample_5","Sample_6","Sample_7","Sample_8","Sample_9","Sample_10","Sample_11","Sample_12","Sample_13","Sample_14","Sample_15","Sample_16","Sample_17","Sample_18","Sample_19","Sample_20","Sample_21","Sample_22") 
phenotype_angus_holstein <- c("2","2","2","2","2","1","1","1","1","1","2","2","2","2","2", "2","2","1","1","1","1","1")
missing_row <- c(0,0,0,0,0,0,0,0,0,0,0,0,0,0,0,0,0,0,0,0,0,0)
sex <- c(2,2,2,2,2,2,2,2,2,2,2,2,2,2,2,2,2,2,2,2,2,2)
fam_file <- data.frame(sample_id,sample_id,missing_row,missing_row,sex,phenotype_angus_holstein)


#write.table(ah_genotypes, "/mnt/storage/lab_folder/heifer_infertility/AI_angus_holstein/assoc_analysis/angus_holstein_genotypes.lgen", col.names = FALSE, row.names = FALSE, sep = "\t" , quote = FALSE)
#write.table(snp_map_genotypes, "/mnt/storage/lab_folder/heifer_infertility/AI_angus_holstein/assoc_analysis/angus_holstein_genotypes.map", col.names = FALSE, row.names = FALSE, sep = "\t" , quote = FALSE)
#write.table(fam_file, "/mnt/storage/lab_folder/heifer_infertility/AI_angus_holstein/assoc_analysis/angus_holstein_genotypes.fam", col.names = FALSE, row.names = FALSE, sep = "\t" , quote = FALSE)
```

##### Quality control and analysis in Plink

```
system("/home/fbiase/bioinfo/plink --lfile /mnt/storage/lab_folder/heifer_infertility/AI_angus_holstein/assoc_analysis/angus_holstein_genotypes --cow  --recode --out /mnt/storage/lab_folder/heifer_infertility/AI_angus_holstein/assoc_analysis/angus_holstein_genotypes_data")

system("/home/fbiase/bioinfo/plink --file /mnt/storage/lab_folder/heifer_infertility/AI_angus_holstein/assoc_analysis/angus_holstein_genotypes_data --cow --make-bed --out /mnt/storage/lab_folder/heifer_infertility/AI_angus_holstein/assoc_analysis/angus_holstein_genotypes_data")

system("/home/fbiase/bioinfo/plink --bfile /mnt/storage/lab_folder/heifer_infertility/AI_angus_holstein/assoc_analysis/angus_holstein_genotypes_data --cow --test-missing --out /mnt/storage/lab_folder/heifer_infertility/AI_angus_holstein/assoc_analysis/angus_holstein_genotypes_data")

system("perl /mnt/storage/lab_folder/heifer_infertility/AI_angus_holstein/assoc_analysis/run-diffmiss-qc.pl  /mnt/storage/lab_folder/heifer_infertility/AI_angus_holstein/assoc_analysis/angus_holstein_genotypes_data > /mnt/storage/lab_folder/heifer_infertility/AI_angus_holstein/assoc_analysis/fail-diffmiss-qc.txt")

system("/home/fbiase/bioinfo/plink --file /mnt/storage/lab_folder/heifer_infertility/AI_angus_holstein/assoc_analysis/angus_holstein_genotypes_data --cow --mind 0.1   --maf 0.01 --geno 0.05 --hwe 0.00001 --make-bed --autosome --out /mnt/storage/lab_folder/heifer_infertility/AI_angus_holstein/assoc_analysis/angus_holstein_genotypes_qc_data")
#606553 variants and 22 cattle pass filters and QC

system("/home/fbiase/bioinfo/plink --bfile /mnt/storage/lab_folder/heifer_infertility/AI_angus_holstein/assoc_analysis/angus_holstein_genotypes_qc_data --cow --recode --tab --out /mnt/storage/lab_folder/heifer_infertility/AI_angus_holstein/assoc_analysis/angus_holstein_genotypes_qc_data")

system("/home/fbiase/bioinfo/plink --bfile /mnt/storage/lab_folder/heifer_infertility/AI_angus_holstein/assoc_analysis/angus_holstein_genotypes_qc_data  --cow --indep-pairwise 50 5 0.2 --out /mnt/storage/lab_folder/heifer_infertility/AI_angus_holstein/assoc_analysis/angus_holstein_genotypes_qc_data")

system("/home/fbiase/bioinfo/plink --bfile /mnt/storage/lab_folder/heifer_infertility/AI_angus_holstein/assoc_analysis/angus_holstein_genotypes_qc_data --cow  --exclude /mnt/storage/lab_folder/heifer_infertility/AI_angus_holstein/assoc_analysis/angus_holstein_genotypes_qc_data.prune.in --pca --out /mnt/storage/lab_folder/heifer_infertility/AI_angus_holstein/assoc_analysis/angus_holstein_genotypes_qc_data")

system("/home/fbiase/bioinfo/plink --bfile /mnt/storage/lab_folder/heifer_infertility/AI_angus_holstein/assoc_analysis/angus_holstein_genotypes_qc_data --cow  --cluster --K 2 --out /mnt/storage/lab_folder/heifer_infertility/AI_angus_holstein/assoc_analysis/angus_holstein_genotypes_qc_data_cluster")

system("/home/fbiase/bioinfo/plink --bfile /mnt/storage/lab_folder/heifer_infertility/AI_angus_holstein/assoc_analysis/angus_holstein_genotypes_qc_data --cow --within /mnt/storage/lab_folder/heifer_infertility/AI_angus_holstein/assoc_analysis/angus_holstein_genotypes_qc_data_cluster.cluster2 --exclude /mnt/storage/lab_folder/heifer_infertility/AI_angus_holstein/assoc_analysis/angus_holstein_genotypes_qc_data.prune.in --assoc fisher --out /mnt/storage/lab_folder/heifer_infertility/AI_angus_holstein/assoc_analysis/angus_holstein_genotypes_qc_data")


angus_holstein_genotypes_qc_data.fisher<-data.table::fread("/mnt/storage/lab_folder/heifer_infertility/AI_angus_holstein/assoc_analysis/angus_holstein_genotypes_qc_data.assoc.fisher", data.table=FALSE)
#head(angus_holstein_genotypes_qc_data.fisher)
head(angus_holstein_genotypes_qc_data.fisher<-angus_holstein_genotypes_qc_data.fisher[order(angus_holstein_genotypes_qc_data.fisher$P),], n=20)
```

##### load the analysis

```
angus_holstein_genotypes_qc_data.fisher<-data.table::fread("/mnt/storage/lab_folder/heifer_infertility/AI_angus_holstein/assoc_analysis/angus_holstein_genotypes_qc_data.assoc.fisher", data.table=FALSE)
angus_holstein_genotypes_qc_data.fisher<-angus_holstein_genotypes_qc_data.fisher<-angus_holstein_genotypes_qc_data.fisher[order(angus_holstein_genotypes_qc_data.fisher$P),]

head(angus_holstein_genotypes_qc_data.fisher)
```

```
##        CHR                    SNP        BP A1    F_A    F_U A2         P
## 319065  12     BovineHD1200026258  89664275  G 0.7083 0.0000  A 6.753e-07
## 282234  11 Hapmap31720-BTA-126418  37519673  G 0.1250 0.7778  A 2.658e-05
## 319054  12     BovineHD1200026243  89607810  G 0.5000 0.0000  A 1.342e-04
## 148145   5     BovineHD0500034893 119441043  G 0.1667 0.7500  A 1.727e-04
## 148146   5     BovineHD0500034894 119441844  G 0.1667 0.7500  T 1.727e-04
## 541682  27     BovineHD2700000503   1491696  C 0.1667 0.7500  T 1.727e-04
##             OR
## 319065      NA
## 282234 0.04082
## 319054      NA
## 148145 0.06667
## 148146 0.06667
## 541682 0.06667
```

```
#angus_holstein_genotypes_qc_data.fisher$label<-""
#angus_holstein_genotypes_qc_data.fisher$label<-ifelse(angus_holstein_genotypes_qc_data.fisher$P <= 2.658e-05, paste("chr",angus_holstein_genotypes_qc_data.fisher$CHR,":", angus_holstein_genotypes_qc_data.fisher$BP, sep="") ,"")
```

##### Genotypes per sample

```
ah_genotypes<- data.table::fread("/mnt/storage/lab_folder/heifer_infertility/AI_angus_holstein/assoc_analysis/Virginia_Tech_Univ_Biase_BOV770V01_20220825_FinalReport.txt", header = TRUE, sep = "\t", skip = 9)
ah_genotypes<-ah_genotypes[,c(1,2,3,4)]
colnames(ah_genotypes)<-c("SNP_Name", "Sample_ID","Allele1", "Allele2")
ah_genotypes<- ah_genotypes[!ah_genotypes$Sample_ID=="Sample_23" & !ah_genotypes$Sample_ID=="Sample_24",]

ah_genotypes<-ah_genotypes[ah_genotypes$SNP_Name %in% angus_holstein_genotypes_qc_data.fisher$SNP,]
ah_genotypes$genotype<-paste(ah_genotypes$Allele1,ah_genotypes$Allele2, sep="")
ah_genotypes<-ah_genotypes[!(ah_genotypes$genotype == "--"),]

ah_genotypes %>% group_by(Sample_ID) %>% summarise(n = n()) %>% print(n = 24)
```

```
## # A tibble: 24 × 2
##    Sample_ID      n
##    <chr>      <int>
##  1 Sample 1  574813
##  2 Sample 10 574778
##  3 Sample 11 574705
##  4 Sample 12 574938
##  5 Sample 13 574757
##  6 Sample 14 574715
##  7 Sample 15 574739
##  8 Sample 16 574945
##  9 Sample 17 574960
## 10 Sample 18 574955
## 11 Sample 19 574816
## 12 Sample 2  574705
## 13 Sample 20 574893
## 14 Sample 21 574915
## 15 Sample 22 574850
## 16 Sample 23 574782
## 17 Sample 24 574817
## 18 Sample 3  574592
## 19 Sample 4  574959
## 20 Sample 5  574882
## 21 Sample 6  574902
## 22 Sample 7  574791
## 23 Sample 8  574849
## 24 Sample 9  574909
```

#### Transcriptome analysis

##### Read count summary

```
files<-list.files("/mnt/storage/lab_folder/heifer_infertility/AI_angus_holstein/alignment3", recursive=T, pattern="summary.txt", full.names = TRUE)
#length(files)

reads_produced_a<-data.frame()
reads_produced_b<-data.frame()
#length(files)

for (n in 1:22) {
reads_produced<-read.delim(files[n], sep= " ",header=FALSE, stringsAsFactors = FALSE, comment.char= "#")
sample<-substring(files[n], 73,74)
number_of_reads_produced<-as.integer(reads_produced[1,1])

reads_produced_a<-data.frame(sample,number_of_reads_produced)
reads_produced_b<-rbind(reads_produced_b,reads_produced_a)
}

files<-list.files("/mnt/storage/lab_folder/heifer_infertility/AI_angus_holstein/counting3", recursive=T, pattern="count.summary", full.names = TRUE)
#length(files)
counting_summary_a<-data.frame()
counting_summary_b<-data.frame()
#length(files)

for (n in 1:22) {
counting_summary<-read.delim(files[n], sep= "\t",header=TRUE, stringsAsFactors = FALSE, comment.char= "#")
sample<-substring(files[n], 72,73)
number_of_reads_assigned<-counting_summary[1,2]
unassigned_NoFeatures<-counting_summary[12,2]
unassigned_Ambiguity<-counting_summary[14,2]
counting_summary_a<-data.frame(sample,number_of_reads_assigned, unassigned_NoFeatures, unassigned_Ambiguity)
counting_summary_b<-rbind(counting_summary_b,counting_summary_a)
}

counting_summary_b$reads_sequenced<-reads_produced_b$number_of_reads_produced
counting_summary_b$reads_retained<-counting_summary_b$number_of_reads_assigned + counting_summary_b$unassigned_NoFeatures + counting_summary_b$unassigned_Ambiguity
counting_summary_b$reads_discarted<-counting_summary_b$reads_sequenced - counting_summary_b$reads_retained
counting_summary_b$perc_number_of_reads_discarted<-counting_summary_b$reads_discarted/counting_summary_b$reads_sequenced
counting_summary_b$perc_number_of_reads_retained<-counting_summary_b$reads_retained/counting_summary_b$reads_sequenced
counting_summary_b$perc_number_of_reads_assigned<-counting_summary_b$number_of_reads_assigned/counting_summary_b$reads_sequenced
counting_summary_b$perc_unassigned_NoFeatures<-counting_summary_b$unassigned_NoFeatures/counting_summary_b$reads_sequenced
counting_summary_b$perc_unassigned_Ambiguity<-counting_summary_b$unassigned_Ambiguity/counting_summary_b$reads_sequenced

#write.table(counting_summary_b, file="/mnt/storage/lab_folder/shared_R_codes/chace/angus_holstein_AI/resources/counting_summary_b_second_alignment.txt",quote = FALSE, sep = "\t",row.names = FALSE)
```

##### Plot read count summary

```
counting_summary_c<-reshape2::melt(counting_summary_b[,c(1,8,10:12)])
```

```
## Using sample as id variables
```

```
counting_summary_c$value<-counting_summary_c$value*100
counting_summary_c$variable<-factor(counting_summary_c$variable, levels=c( "perc_number_of_reads_discarted" ,"perc_unassigned_Ambiguity","perc_unassigned_NoFeatures" , "perc_number_of_reads_assigned"))
plot_a<-ggplot() + 
  geom_bar(aes(y = value, x = sample, fill = variable), data = counting_summary_c,stat="identity")+
  scale_y_continuous(name="Percentage", breaks = seq(0,100,10))+
  scale_fill_hue(labels=c("Discarted", "Unassigned_Ambiguity","Unassigned_NoFeatures" , "Assigned"))+
  #ggtitle("Distribution of reads after alignment")+
  theme_bw(base_size = 12)+
  theme(legend.title = element_blank(),
        axis.title.x = element_blank(),
        axis.text.x = element_text(angle=75, hjust=1))

counting_summary_c<-reshape2::melt(counting_summary_b[,c(1,5)])
```

```
## Using sample as id variables
```

```
plot_b<-ggplot() + 
  geom_bar(aes(y = value, x = sample, fill = variable), data = counting_summary_c,stat="identity")+
  scale_y_continuous(name="Read pairs sequenced")+
  theme_bw(base_size = 12)+
  theme(legend.title = element_blank(),
        axis.title.x = element_blank(),
        axis.text.x = element_blank())

counting_summary_c<-reshape2::melt(counting_summary_b[,c(1,2)])
```

```
## Using sample as id variables
```

```
plot_c<-ggplot() + 
  geom_bar(aes(y = value, x = sample, fill = variable), data = counting_summary_c,stat="identity")+
  scale_y_continuous(name="Read pairs in annotation")+
  geom_hline(yintercept=10^7, linetype="dashed", color = "gray", size=1)+
  theme_bw(base_size = 12)+
  theme(legend.title = element_blank(),
        axis.title.x = element_blank(),
        axis.text.x = element_blank())
```

```
## Warning: Using `size` aesthetic for lines was deprecated in ggplot2 3.4.0.
## ℹ Please use `linewidth` instead.
```

```
counting_summary_c<-reshape2::melt(counting_summary_b[,c(1,10)])
```

```
## Using sample as id variables
```

```
plot_d<-ggplot() + 
  geom_bar(aes(y = value, x = sample, fill = variable), data = counting_summary_c,stat="identity")+
  scale_y_continuous(name="Proportion reads \n matching annotation")+
  theme_bw(base_size = 12)+
  theme(legend.title = element_blank(),
        axis.title.x = element_blank(),
        axis.text.x = element_blank())

plot_grid( plot_b, plot_c,plot_d, plot_a,  ncol = 1, align = 'v')
```

##### Import Sample Information

```
sample_info <- as.data.frame(read_excel("/mnt/storage/lab_folder/shared_R_codes/chace/angus_holstein_AI/resources/sample_info.xlsx"))
sample_info$sample<- as.factor(sample_info$sample)
sample_info$preg<- as.factor(sample_info$preg)
sample_info$breed<- as.factor(sample_info$breed)
```

##### Counts per sample

```
files<-list.files("/mnt/storage/lab_folder/heifer_infertility/AI_angus_holstein/counting3", recursive=T, pattern="count", full.names = TRUE)
files<-files[grep("summary", files, invert = TRUE)]
files<-files[grep(".count1", files, invert = TRUE)]
files<-files[grep(".count2", files, invert = TRUE)]
files<-files[grep(".sh", files, invert = TRUE)]
#length(files)
count_data<-data.frame(matrix(nrow=27607))
for (n in 1:22) {
  count<-read.delim(files[n], header=TRUE, sep= "\t", stringsAsFactors = FALSE, comment.char= "#")
  count<-count[,c(1,7)]
  count_data<-cbind(count_data,count)
}
rownames(count_data)<-count_data[,2]
count_data<-count_data[,seq(from = 3, to = 45, by = 2)]
colnames(count_data)<- substr(colnames(count_data), 74, 75)

colnames(count_data)<-sample_info$Animal_ID
#colnames(count_data)<-sample_info$SNP_array_ID
#count_data_export<-data.frame(gene_id=rownames(count_data),count_data)
#write_delim(count_data_export, file = "/mnt/storage/lab_folder/shared_R_codes/fernando/angus_holstein_association/resources/2022_12_02_unfiltered_count_data.txt", delim = "\t", quote =  "none")
#system("bzip2 /mnt/storage/lab_folder/shared_R_codes/fernando/angus_holstein_association/resources/2022_12_02_unfiltered_count_data.txt")
```

##### Sequencing summary

```
count_data_a<-count_data[rowSums(count_data)>0,]
count_data_annotated<-merge(count_data_a,annotation.ensembl.symbol, by.x="row.names", by.y="ensembl_gene_id", all.x=TRUE, all.y=FALSE)

n_reads_protein_coding<-sum(count_data_annotated[count_data_annotated$gene_biotype=='protein_coding',c(2:6)])
n_reads_lncRNA<-sum(count_data_annotated[count_data_annotated$gene_biotype=='lncRNA',c(2:6)])
n_reads_pseudogene<-sum(count_data_annotated[count_data_annotated$gene_biotype %in% c('pseudogene','processed_pseudogene'),c(2:6)])
n_reads_others<-sum(count_data_annotated[!(count_data_annotated$gene_biotype %in% c('pseudogene','processed_pseudogene','protein_coding','lncRNA')),c(2:6)])

summary_RNA_seq<-data.frame(class=c('protein_coding','lncRNA', 'pseudogene','others'), nreads=c(n_reads_protein_coding,n_reads_lncRNA,n_reads_pseudogene,n_reads_others))

summary_RNA_seq<-mutate(summary_RNA_seq,  prop = round(nreads/sum(nreads)*100,2))
summary_RNA_seq<-summary_RNA_seq[with(summary_RNA_seq, order(-nreads)),]
summary_RNA_seq$class<-factor(summary_RNA_seq$class, levels=c('protein_coding', 'lncRNA','pseudogene','others' ))

knitr::kable(summary_RNA_seq, format = 'pandoc')
```

|  | class | nreads | prop |
| --- | --- | --- | --- |
| 1 | protein\_coding | 123653221 | 98.85 |
| 4 | others | 570242 | 0.46 |
| 3 | pseudogene | 442795 | 0.35 |
| 2 | lncRNA | 422486 | 0.34 |

##### Subset for protein-coding, lncRNAs, and pseudogenes

```
count_data_a<-count_data[rowSums(count_data)>0,]
count_data_annotated<-merge(count_data_a,annotation.ensembl.symbol, by.x="row.names", by.y="ensembl_gene_id", all.x=TRUE, all.y=FALSE)
count_data_annotated<-count_data_annotated[count_data_annotated$gene_biotype %in% c('protein_coding', 'lncRNA','pseudogene'),]
count_data_annotated_length<-gene.length[gene.length$ensembl_gene_id %in% count_data_annotated$Row.names, 2]
```

##### Calculate CPM, FPKM, TPM

```
count_data_b<-count_data_annotated[,c(2:23)]
rownames(count_data_b)<-count_data_annotated$Row.names

lib_size <- base::colSums(count_data_b)
norm_factors <- edgeR::calcNormFactors(object = count_data_b, lib.size = lib_size, method = "TMM")

#FPKM
data_fpkm<-edgeR::rpkm(sweep(count_data_b, 2, norm_factors, "/"), count_data_annotated_length)

#CPM
data_cpm<-edgeR::cpm(sweep(count_data_b, 2, norm_factors, "/"),normalized.lib.sizes = TRUE,log = FALSE)

#TPM
x <- count_data_b / count_data_annotated_length
data_tpm <- data.frame(t( t(x) * 1e6 / colSums(x) ))
rownames(data_tpm)<-count_data_annotated$Row.names

keep<-rowSums(data_fpkm>1) >=5
data_fpkm_filtered<-data_fpkm[keep,]
keep<-rowSums(data_cpm>1) >=5
data_cpm_filtered<-data_cpm[keep,]
genes_expressed<-intersect(rownames(data_fpkm_filtered), rownames(data_cpm_filtered))

data_tpm<-data_tpm[rownames(data_tpm) %in% genes_expressed,]

data_fpkm_filtered<-data_fpkm_filtered[rownames(data_fpkm_filtered) %in% genes_expressed,]
data_cpm_filtered<-data_cpm_filtered[rownames(data_cpm_filtered) %in% genes_expressed,]

count_data_filtered<-count_data[rownames(count_data) %in% genes_expressed,]
```

##### Gene summary

```
table(annotation.ensembl.symbol[annotation.ensembl.symbol$ensembl_gene_id %in% genes_expressed, ]$gene_biotype)
```

```
## 
##         lncRNA protein_coding     pseudogene 
##            228          12105            112
```

##### DEG Analysis - Fertility

```
heifer_data <- sample_info

#edgeR
preg<-factor(heifer_data$preg, levels=c("P", "NP"))
breed<-factor(heifer_data$breed, levels=c("ANG","HOL"))
heifer_design<- model.matrix(~ heifer_data$breed + heifer_data$preg)

heifer <-edgeR::DGEList(count=count_data_filtered, group=heifer_data$preg)
heifer <-edgeR::estimateDisp(heifer, heifer_design, robust=TRUE)
heifer_QLFit <- edgeR::glmQLFit(heifer, heifer_design, robust=TRUE)
heifer_QLF <- edgeR::glmQLFTest(heifer_QLFit, coef="heifer_data$pregP")
heifer_edgeR_results_QLF<- edgeR::topTags(heifer_QLF, adjust.method = "fdr", n=Inf)$table

#DESeq2 wald
heifer_DESeq <- DESeq2::DESeqDataSetFromMatrix(countData=count_data_filtered, colData=heifer_data, design= ~ breed + preg)
heifer_Wald<-DESeq2::DESeq(heifer_DESeq, test="Wald")
heifer_DESeq_results_Wald<-DESeq2::results(heifer_Wald, contrast=c("preg",  "P", "NP"), pAdjustMethod="fdr", tidy=TRUE)

#DESeq2 LRT
heifer_LRT<-DESeq2::DESeq(heifer_DESeq, test="LRT", full= ~ breed + preg, reduced= ~ breed )
heifer_DESeq_results_LRT<-DESeq2::results(heifer_LRT, contrast=c("preg",  "P", "NP"), pAdjustMethod="fdr", tidy=TRUE)


heifer_edgeR_results_QLF_heifer_DESeq_results_Wald_LRT<-merge(heifer_edgeR_results_QLF,heifer_DESeq_results_Wald, by.x="row.names", by.y="row")
heifer_edgeR_results_QLF_heifer_DESeq_results_Wald_LRT<-merge(heifer_edgeR_results_QLF_heifer_DESeq_results_Wald_LRT, heifer_DESeq_results_LRT, by.x="Row.names",by.y="row")

heifer_edgeR_results_QLF_heifer_DESeq_results_Wald_LRT<-heifer_edgeR_results_QLF_heifer_DESeq_results_Wald_LRT[with(heifer_edgeR_results_QLF_heifer_DESeq_results_Wald_LRT, order(PValue)),]


heifer_edgeR_results_QLF_heifer_DESeq_results_Wald_LRT[heifer_edgeR_results_QLF_heifer_DESeq_results_Wald_LRT$PValue < 0.002 & heifer_edgeR_results_QLF_heifer_DESeq_results_Wald_LRT$pvalue.x< 0.001 & heifer_edgeR_results_QLF_heifer_DESeq_results_Wald_LRT$pvalue.y < 0.001,  ]
```

```
##               Row.names      logFC   logCPM        F       PValue       FDR
## 1811 ENSBTAG00000004278  0.2149330 5.622078 15.39755 0.0007064172 0.9997511
## 1274 ENSBTAG00000002972 -0.2948188 2.240588 12.99729 0.0015381391 0.9997511
##      baseMean.x log2FoldChange.x    lfcSE.x    stat.x     pvalue.x    padj.x
## 1811  1169.0159        0.2122938 0.05272878  4.026147 5.669837e-05 0.7055545
## 1274   110.8074       -0.2984712 0.08631589 -3.457894 5.444147e-04 0.9998599
##      baseMean.y log2FoldChange.y    lfcSE.y   stat.y     pvalue.y    padj.y
## 1811  1169.0159        0.2122938 0.05272878 16.18059 5.758103e-05 0.7165383
## 1274   110.8074       -0.2984712 0.08631589 11.96050 5.434015e-04 0.9998482
```

```
#heifer_edgeR_results_QLF_heifer_DESeq_results_Wald_LRT_annotated<-merge(heifer_edgeR_results_QLF_heifer_DESeq_results_Wald_LRT, annotation.ensembl.symbol, by.x="Row.names", by.y="ensembl_gene_id", all.x=TRUE, all.y=FALSE)

#write.table(heifer_edgeR_results_QLF_heifer_DESeq_results_Wald_LRT_annotated, "/mnt/storage/lab_folder/shared_R_codes/fernando/angus_holstein_association/results/Additional_file_3.txt",quote = FALSE, sep = "\t",row.names = FALSE, col.names = TRUE)
```

##### Calculate q-value - Fertility

```
#Create the permutations matrix
perm_without_replacement <- function(n, r){
 return(factorial(n)/factorial(n - r))
}

permutation_matrix<-matrix(nrow=10000,ncol=22)
#head(permutation_matrix)
dim(permutation_matrix)

for (i in seq(1:10000)){
 sampling <- sample(1:22,22, replace = FALSE)
 if ( ! identical(sampling , c(1:22))){
 permutation_matrix[i,]<-sample(1:22,22, replace = FALSE)
 }}

permutation_matrix<-permutation_matrix[!duplicated(permutation_matrix),]
dim(permutation_matrix)
#head(permutation_matrix)

#permutation_matrix<-permutation_matrix[sample(1:dim(permutation_matrix)[1], 100),]
rand<-dim(permutation_matrix)[1]

sequence.pvalue<-seq(0.0005, 0.002, 0.0001)

#edgeR
results <- filebacked.big.matrix(length(sequence.pvalue),rand, type="double", init=0, separated=FALSE,
 backingfile="incidence_matrix.bin",
 descriptor="incidence_matrix.desc")
mdesc_result<- describe(results)

cl <- makeCluster(30)
registerDoParallel(cl)
results[,]<-foreach(i = sequence.pvalue, .combine='rbind', .inorder=TRUE, .packages=c("edgeR","bigmemory"), .verbose=FALSE) %:%

 foreach(j = 1:rand, .combine='cbind', .inorder=FALSE,.packages=c("edgeR","bigmemory"), .verbose=FALSE ) %dopar%
{

 preg<-factor(heifer_data$preg, levels=c("P", "NP"))
 breed<-factor(heifer_data$breed, levels=c("ANG","HOL"))

 design <- model.matrix(~ breed + preg)
 dds<-DGEList(count=count_data_filtered[,permutation_matrix[j,]], group=heifer_data$preg)
 dds<-estimateDisp(dds, design, robust=TRUE)
 dds <- glmFit(dds, design)
 dds <- glmQLFTest(heifer_QLFit, coef=3)
 length(which(topTags(dds,n=Inf)$table$PValue < i))

 }
stopCluster(cl)
total.rand <- rand * 12065
qvalue<-data.frame(raw.pvalue = sequence.pvalue,
 e.pvalue.edgeR= (rowSums(results[,]+1))/(total.rand+1),
 e.pvalue.edgeR.round= round((rowSums(results[,]+1))/(total.rand+1) ,4))

#qvalue
rm(results)
system("rm incidence_matrix.bin")
system("rm incidence_matrix.desc")

#DESEQ2
results.rand <- filebacked.big.matrix(length(sequence.pvalue),rand, type="double", init=0, separated=FALSE,
 backingfile="incidence_matrix.bin",
 descriptor="incidence_matrix.desc")
mdesc_result<- describe(results.rand)
cl <- makeCluster(30)
registerDoParallel(cl)
results.rand[,]<-foreach(i = sequence.pvalue, .combine='rbind', .inorder=TRUE, .packages=c("DESeq2","bigmemory"), .verbose=FALSE) %:%

 foreach(j = 1:rand, .combine='cbind', .inorder=FALSE,.packages=c("DESeq2","bigmemory"), .verbose=TRUE ) %dopar%
{

heifer_DESeq <-DESeqDataSetFromMatrix(countData=count_data_filtered[permutation_matrix[j,]], colData=heifer_data, design= ~   breed + preg)
heifer_Wald<-DESeq(heifer_DESeq, test="Wald")
heifer_DESeq_results_Wald<-results(heifer_Wald, contrast=c("preg",  "P", "NP"), pAdjustMethod="fdr", tidy=TRUE)

length(which(heifer_DESeq_results_Wald$pvalue < i))

 }
stopCluster(cl)
#results.rand[1:5,1:5]
total.rand <- rand * 12065

qvalue2<-data.frame(raw.pvalue = sequence.pvalue,
 e.pvalue.DESEQ2= (rowSums(results.rand[,]+1))/(total.rand+1),
 e.pvalue.DESEQ2.round= round((rowSums(results.rand[,]+1))/(total.rand+1) ,4))

rm(results.rand)
system("rm incidence_matrix.bin")
system("rm incidence_matrix.desc")


#DESEQ2
results.rand <- filebacked.big.matrix(length(sequence.pvalue),rand, type="double", init=0, separated=FALSE,
 backingfile="incidence_matrix.bin",
 descriptor="incidence_matrix.desc")
mdesc_result<- describe(results.rand)
cl <- makeCluster(30)
registerDoParallel(cl)
results.rand[,]<-foreach(i = sequence.pvalue, .combine='rbind', .inorder=TRUE, .packages=c("DESeq2","bigmemory"), .verbose=FALSE) %:%

 foreach(j = 1:rand, .combine='cbind', .inorder=FALSE,.packages=c("DESeq2","bigmemory"), .verbose=TRUE ) %dopar%
{

heifer_DESeq <-DESeqDataSetFromMatrix(countData=count_data_filtered[permutation_matrix[j,]], colData=heifer_data, design= ~  breed + preg)
heifer_LRT<-DESeq2::DESeq(heifer_DESeq, test="LRT", full= ~ breed + preg, reduced= ~ breed )
heifer_DESeq_results_LRT<-DESeq2::results(heifer_LRT, contrast=c("preg",  "P", "NP"), pAdjustMethod="fdr", tidy=TRUE)

length(which(heifer_DESeq_results_LRT$pvalue < i))

 }
stopCluster(cl)
#results.rand[1:5,1:5]
total.rand <- rand * 12065

qvalue3<-data.frame(raw.pvalue = sequence.pvalue,
 e.pvalue.DESEQ2= (rowSums(results.rand[,]+1))/(total.rand+1),
 e.pvalue.DESEQ2.round= round((rowSums(results.rand[,]+1))/(total.rand+1) ,4))
#qvalue2
qvalue4<-cbind(qvalue,qvalue2,qvalue3)
#qvalue4

rm(results.rand)
system("rm incidence_matrix.bin")
system("rm incidence_matrix.desc")

write.table(qvalue4,file="/mnt/storage/lab_folder/shared_R_codes/fernando/angus_holstein_association/results/2022_10_09_efdr.txt", sep = "\t",append = FALSE, quote = FALSE)
```

#### Proteome analysis

##### Load and process the data

```
holstein_data<- data.frame(read_excel( "/mnt/storage/lab_folder/shared_R_codes/fernando/angus_holstein_association/resources/220915_Bovine_BS1_01.xlsx"))
angus_data<- data.frame(read_excel( "/mnt/storage/lab_folder/shared_R_codes/fernando/angus_holstein_association/resources/221101_Bovine_Serum.xlsx"))

holstein_data<-holstein_data[holstein_data$Accession %in% angus_data$Accession,]
angus_data<- angus_data[angus_data$Accession %in% holstein_data$Accession,]

proteome_data<-cbind(holstein_data, angus_data)

row.names(proteome_data)<-proteome_data$Accession
#colnames(proteome_data[, c(38:57, 135:158)])
proteome_data<-proteome_data[, c(38:57, 135:158)]
#colnames(proteome_data)

#head(proteome_data)

proteome_data<- proteome_data[rowSums(is.na(proteome_data))<1,]
proteome_data[is.na(proteome_data)]<-0
proteome_data<-log(proteome_data+0.1, base=2)
```

##### Analysis with generalized mixed models

```
heifer_data   <- as.data.frame(read_excel("/mnt/storage/lab_folder/shared_R_codes/chace/angus_holstein_AI/resources/sample_info.xlsx"))
preg_factor   <- rep(as.factor(heifer_data$preg), each=2)
breed_factor  <- rep(as.factor(heifer_data$breed), each=2)
subject_factor<-factor(rep(c("A","B","C", "D", "E", "F", "G", "H", "I", "J", "K", "L", "M", "N", "O", "P", "Q", "R", "S", "T", "U", "V"),each=2))
```

```
data_frame_results<-data.frame()

for(i in 1:dim(proteome_data)[1]){
  data_frame_aov<- data.frame(values = t(proteome_data[i,])[,1],groups=preg_factor, breed_factor, subject=subject_factor, row.names = NULL)
  model<-lmer(values ~ groups + breed_factor + (1|subject), data=data_frame_aov)
  anova_results<-Anova(model, type="III", test.statistic="F")
  estimate<-summary(pairs(emmeans(model, "groups", data=data_frame_aov), adjust="none"))
  data_frame_results<-rbind(data_frame_results, data.frame(protein=rownames(proteome_data[i,]),F_stat=anova_results$F[2], p_value=anova_results$`Pr(>F)`[2] ,
                            contrast= estimate$contrast, estimate=estimate$estimate, SE=estimate$SE, p_value_t=estimate$p.value))
}

data_frame_results$fdr<-p.adjust(data_frame_results$p_value, method="fdr")
```

##### Analysis with LIMMA

```
design <- model.matrix(~ preg_factor + breed_factor)
dupcor <- duplicateCorrelation(proteome_data,design,block=subject_factor)

fit <- lmFit(proteome_data, design, block=subject_factor,correlation=dupcor$consensus,method="robust")
fit1 <- eBayes(fit)
results_limma<-topTable(fit1,coef=2,adjust.method="fdr", number=Inf)
```

##### Analysis with LIMMA

```
combined_results<-merge(data_frame_results, results_limma, by.x="protein", by.y='row.names')
#combined_results[combined_results$fdr<0.05 & combined_results$adj.P.Val <0.05,]
combined_results<-combined_results[order(combined_results$p_value),]
head(combined_results)
```

```
##     protein    F_stat      p_value contrast   estimate        SE    p_value_t
## 79   A5D798 26.389847 5.858447e-05   NP - P -0.5572665 0.1084787 5.858447e-05
## 159  P19034 15.839204 8.022836e-04   NP - P  1.3674751 0.3435997 8.022836e-04
## 117  F1MYX5 15.778840 8.163165e-04   NP - P  1.0870542 0.2736615 8.163165e-04
## 151  P02768  9.470759 6.198639e-03   NP - P  0.6402478 0.2080443 6.198639e-03
## 120  F1N102  9.407474 6.342328e-03   NP - P  0.6878992 0.2242788 6.342328e-03
## 124  G3MY71  9.227813 6.770946e-03   NP - P -1.0526458 0.3465236 6.770946e-03
##            fdr      logFC  AveExpr         t      P.Value    adj.P.Val
## 79  0.01247849  0.5797261 32.11571  8.029803 3.996312e-10 8.512144e-08
## 159 0.05795847 -1.2511109 24.79571 -5.911072 4.755260e-07 5.064352e-05
## 117 0.05795847 -1.0391210 25.41215 -5.310643 3.545060e-06 1.887745e-04
## 151 0.24036858 -0.6760556 31.26988 -5.018421 9.319662e-06 3.970176e-04
## 120 0.24036858 -0.7003252 30.61259 -5.328767 3.337782e-06 1.887745e-04
## 124 0.24036858  1.0083131 27.69917  3.809532 4.335290e-04 7.695139e-03
##              B
## 79  12.9085204
## 159  5.9190017
## 117  3.9442552
## 151  2.9836050
## 120  3.9869840
## 124 -0.7425267
```

```
#write.table(combined_results, "/mnt/storage/lab_folder/shared_R_codes/fernando/angus_holstein_association/results/Additional_file_4.txt",quote = FALSE, sep = "\t",row.names = FALSE, col.names = TRUE)
```

##### Obtain breed specific estimates

###### A5D798 Alpha-ketoglutarate-dependent dioxygenase FTO

```
  data_frame_aov<- data.frame(values = t(proteome_data[rownames(proteome_data)=="A5D798",])[,1],groups=preg_factor, breed_factor, subject=subject_factor, row.names = NULL)
  model<-lmer(values ~ groups*breed_factor + (1|subject), data=data_frame_aov)
  anova_results<-Anova(model, type="III", test.statistic="F")
  model %>% emmeans(pairwise ~ groups | breed_factor)
```

```
## $emmeans
## breed_factor = ANG:
##  groups emmean    SE df lower.CL upper.CL
##  NP       31.2 0.091 18     31.0     31.3
##  P        31.5 0.108 18     31.3     31.8
## 
## breed_factor = HOL:
##  groups emmean    SE df lower.CL upper.CL
##  NP       32.7 0.108 18     32.5     32.9
##  P        33.4 0.108 18     33.2     33.7
## 
## Degrees-of-freedom method: kenward-roger 
## Confidence level used: 0.95 
## 
## $contrasts
## breed_factor = ANG:
##  contrast estimate    SE df t.ratio p.value
##  NP - P     -0.395 0.141 18  -2.798  0.0119
## 
## breed_factor = HOL:
##  contrast estimate    SE df t.ratio p.value
##  NP - P     -0.747 0.152 18  -4.906  0.0001
## 
## Degrees-of-freedom method: kenward-roger
```

###### P19034 Apolipoprotein C-II

```
  data_frame_aov<- data.frame(values = t(proteome_data[rownames(proteome_data)=="P19034",])[,1],groups=preg_factor, breed_factor, subject=subject_factor, row.names = NULL)
  model<-lmer(values ~ groups*breed_factor + (1|subject), data=data_frame_aov)
  anova_results<-Anova(model, type="III", test.statistic="F")
  model %>% emmeans(pairwise ~ groups | breed_factor)
```

```
## $emmeans
## breed_factor = ANG:
##  groups emmean    SE df lower.CL upper.CL
##  NP       24.2 0.250 18     23.7     24.7
##  P        23.7 0.296 18     23.0     24.3
## 
## breed_factor = HOL:
##  groups emmean    SE df lower.CL upper.CL
##  NP       26.9 0.296 18     26.3     27.6
##  P        24.6 0.296 18     24.0     25.2
## 
## Degrees-of-freedom method: kenward-roger 
## Confidence level used: 0.95 
## 
## $contrasts
## breed_factor = ANG:
##  contrast estimate    SE df t.ratio p.value
##  NP - P      0.546 0.388 18   1.410  0.1757
## 
## breed_factor = HOL:
##  contrast estimate    SE df t.ratio p.value
##  NP - P      2.325 0.419 18   5.553  <.0001
## 
## Degrees-of-freedom method: kenward-roger
```

###### F1MYX5 Lymphocyte cytosolic protein 1

```
  data_frame_aov<- data.frame(values = t(proteome_data[rownames(proteome_data)=="F1MYX5",])[,1],groups=preg_factor, breed_factor, subject=subject_factor, row.names = NULL)
  model<-lmer(values ~ groups*breed_factor + (1|subject), data=data_frame_aov)
  anova_results<-Anova(model, type="III", test.statistic="F")
  model %>% emmeans(pairwise ~ groups | breed_factor)
```

```
## $emmeans
## breed_factor = ANG:
##  groups emmean    SE df lower.CL upper.CL
##  NP       27.2 0.218 18     26.7     27.6
##  P        26.7 0.258 18     26.2     27.3
## 
## breed_factor = HOL:
##  groups emmean    SE df lower.CL upper.CL
##  NP       24.5 0.258 18     23.9     25.0
##  P        22.6 0.258 18     22.1     23.1
## 
## Degrees-of-freedom method: kenward-roger 
## Confidence level used: 0.95 
## 
## $contrasts
## breed_factor = ANG:
##  contrast estimate    SE df t.ratio p.value
##  NP - P      0.429 0.337 18   1.270  0.2202
## 
## breed_factor = HOL:
##  contrast estimate    SE df t.ratio p.value
##  NP - P      1.855 0.365 18   5.089  0.0001
## 
## Degrees-of-freedom method: kenward-roger
```

#### Multi-omics analysis

##### Preparation of the data for analysis

```
vst_DESeq <- DESeq2::vst( DESeq2::DESeqDataSetFromMatrix(countData=count_data_filtered, colData=sample_info, design= ~ preg + breed), blind = TRUE, nsub = 5000)
vst_DESeq_data<-assay(vst_DESeq)
colnames(vst_DESeq_data)<-heifer_data$SNP_array_ID


vst_DESeq_data<-vst_DESeq_data[rownames(vst_DESeq_data) %in%
heifer_edgeR_results_QLF_heifer_DESeq_results_Wald_LRT[heifer_edgeR_results_QLF_heifer_DESeq_results_Wald_LRT$PValue < 0.01 & heifer_edgeR_results_QLF_heifer_DESeq_results_Wald_LRT$pvalue.x< 0.01 & heifer_edgeR_results_QLF_heifer_DESeq_results_Wald_LRT$pvalue.y < 0.01,  1] ,]

colnames(proteome_data)<-rep(heifer_data$SNP_array_ID, each=2)
proteome_data_mofa<-proteome_data
proteome_data_mofa$proteins<-rownames(proteome_data_mofa)
proteome_data_mofa<-reshape2::melt(proteome_data_mofa)
proteome_data_mofa<-data.frame(proteome_data_mofa %>% group_by(proteins, variable) %>% dplyr::summarize(Mean = mean(value, na.rm=TRUE)), stringsAsFactors = FALSE)
proteome_data_mofa$variable<-as.character(proteome_data_mofa$variable)
proteome_data_mofa<-proteome_data_mofa[!duplicated(proteome_data_mofa[,2:3]),]
proteome_data_mofa<-reshape2::dcast(proteome_data_mofa, proteins ~ variable, value.var ="Mean")
rownames(proteome_data_mofa)<-proteome_data_mofa$proteins
proteome_data_mofa<-proteome_data_mofa[,heifer_data$SNP_array_ID]

proteome_data_mofa<-proteome_data_mofa[rownames(proteome_data_mofa) %in% combined_results[combined_results$p_value<0.05 & combined_results$P.Value<0.05,1],]

holstein_data<- data.frame(read_excel( "/mnt/storage/lab_folder/shared_R_codes/fernando/angus_holstein_association/resources/220915_Bovine_BS1_01.xlsx"))
angus_data<- data.frame(read_excel( "/mnt/storage/lab_folder/shared_R_codes/fernando/angus_holstein_association/resources/221101_Bovine_Serum.xlsx"))
holstein_data<-holstein_data[holstein_data$Accession %in% angus_data$Accession,]
angus_data<- angus_data[angus_data$Accession %in% holstein_data$Accession,]
proteome_data_annotation<-cbind(holstein_data, angus_data)
proteome_data_annotation<-proteome_data_annotation[, c("Accession", "Gene.Symbol")]

proteome_data_mofa<-merge(proteome_data_mofa,proteome_data_annotation, by.x="row.names", by.y="Accession")


ah_genotypes<-data.table::fread("/mnt/storage/lab_folder/heifer_infertility/AI_angus_holstein/assoc_analysis/Virginia_Tech_Univ_Biase_BOV770V01_20220825_FinalReport.txt", header = TRUE, sep = "\t", skip = 9, check.names=TRUE)

ah_genotypes<-ah_genotypes[ah_genotypes$SNP.Name %in% angus_holstein_genotypes_qc_data.fisher[angus_holstein_genotypes_qc_data.fisher$P<0.001,2],]

angus_holstein_genotypes<- ah_genotypes[,c(1,2,7,8 )]
angus_holstein_genotypes$genotype <- paste(angus_holstein_genotypes$Allele1...AB, angus_holstein_genotypes$Allele2...AB, sep="")

angus_holstein_genotypes<- angus_holstein_genotypes[,-c(3,4)]

#head(angus_holstein_genotypes)
angus_holstein_genotypes<-angus_holstein_genotypes[!(angus_holstein_genotypes$Sample.ID %in% c("Sample 23", "Sample 24"))]

angus_holstein_genotypes_a<- as.data.frame(tidyr::spread(angus_holstein_genotypes, Sample.ID, genotype))

rownames(angus_holstein_genotypes_a)<- angus_holstein_genotypes_a$SNP.Name
angus_holstein_genotypes_a<- angus_holstein_genotypes_a[,-1]

colnames(angus_holstein_genotypes_a)<-make.names(colnames(angus_holstein_genotypes_a))

angus_holstein_genotypes_a[angus_holstein_genotypes_a=="AA"]<-as.numeric(0)
angus_holstein_genotypes_a[angus_holstein_genotypes_a=="AB"]<-as.numeric(1)
angus_holstein_genotypes_a[angus_holstein_genotypes_a=="BB"]<-as.numeric(1)

angus_holstein_genotypes_b<-angus_holstein_genotypes_a[rowSums(angus_holstein_genotypes_a=='1',na.rm=TRUE)>=4,]
angus_holstein_genotypes_b<-angus_holstein_genotypes_a[rowSums(angus_holstein_genotypes_a=='2',na.rm=TRUE)>=4,]
angus_holstein_genotypes_b<-angus_holstein_genotypes_a[rowSums(angus_holstein_genotypes_a=='0',na.rm=TRUE)>=4,]

angus_holstein_genotypes_b1<-data.frame(lapply(angus_holstein_genotypes_b,as.numeric))
rownames(angus_holstein_genotypes_b1)<- rownames(angus_holstein_genotypes_b)

angus_holstein_genotypes_b1<-as.matrix(angus_holstein_genotypes_b1,rownames = TRUE)

colnames(angus_holstein_genotypes_b1)<-tolower(gsub(".", "_", colnames(angus_holstein_genotypes_b1),fixed = TRUE))

genotypes_b <- angus_holstein_genotypes_b1[,heifer_data$SNP_array_ID]

snp_map_genotypes <- read.table("/mnt/storage/lab_folder/heifer_infertility/AI_angus_holstein/assoc_analysis/SNP_Map.txt", header = TRUE, sep = "\t")
snp_map_genotypes <- snp_map_genotypes[,c(3,2,4)]

snp_map_genotypes<- snp_map_genotypes[!snp_map_genotypes$Chromosome=="0" & !snp_map_genotypes$Chromosome=="X" & !snp_map_genotypes$Chromosome=="Y",]

genotypes_b<-genotypes_b[rownames(genotypes_b) %in% snp_map_genotypes$Name,]

mofa_list_matrix<-list(mRNA=as.matrix(vst_DESeq_data), protein=as.matrix(proteome_data_mofa), genotypes=as.matrix(genotypes_b))

covariates_mofa<-data.frame(row.names=heifer_data$SNP_array_ID, breed=heifer_data$breed, pregnancy=heifer_data$preg)
covariates_mofa$pregnancy<-ifelse(covariates_mofa$pregnancy == "P", "Fertile", "Sub-fertile")
covariates_mofa$breed<-ifelse(covariates_mofa$breed == "HOL", "Holstein", "Angus")
```

##### Organize objects and run analysis

```
multiomics_mofa <- MultiAssayExperiment(
  experiments = mofa_list_matrix, 
  colData = covariates_mofa
)

MOFAobject <- create_mofa_from_MultiAssayExperiment(multiomics_mofa,extract_metadata = TRUE, groups = "breed")

data_opts <- get_default_data_options(MOFAobject)
head(data_opts)
model_opts <- get_default_model_options(MOFAobject)
head(model_opts)
model_opts$likelihoods<-c("gaussian" ,"gaussian", "bernoulli" )
head(model_opts)
train_opts <- get_default_training_options(MOFAobject)
head(train_opts)
train_opts$convergence_mode<-"slow"

set.seed(12345)

MOFAobject <- prepare_mofa(
  object = MOFAobject,
  data_options = data_opts,
  model_options = model_opts,
  training_options = train_opts
)

MOFAobject.trained.group <- run_mofa(MOFAobject, outfile = "/mnt/storage/lab_folder/shared_R_codes/fernando/angus_holstein_association/results/MOFAobject.trained_2022_11_28_group.hdf5", save_data = TRUE, use_basilisk = TRUE)
```

##### Load the model

```
MOFAobject.trained.group<-load_model("/mnt/storage/lab_folder/shared_R_codes/fernando/angus_holstein_association/results/MOFAobject.trained_2022_11_28_group.hdf5", remove_inactive_factors = TRUE)

head(MOFAobject.trained.group@cache$variance_explained$r2_total[[1]])
```

```
##      mRNA   protein genotypes 
##  44.12020  16.55246  70.13337
```

```
head(MOFAobject.trained.group@cache$variance_explained$r2_per_factor[[1]], n=10)
```

```
##                 mRNA     protein    genotypes
## Factor1  0.005228172  0.02095313 64.481634390
## Factor2 44.102950596  3.60692713  0.008542051
## Factor3  0.006075510 12.91058002  0.001576960
## Factor4  0.005947563  0.01400409  5.641619256
```

#### Figures

##### Figure 1

```
evec <- data.table::fread("/mnt/storage/lab_folder/heifer_infertility/AI_angus_holstein/assoc_analysis/angus_holstein_genotypes_qc_data.eigenvec", data.table = FALSE)
eval <- data.table::fread("/mnt/storage/lab_folder/heifer_infertility/AI_angus_holstein/assoc_analysis/angus_holstein_genotypes_qc_data.eigenval", data.table = FALSE)

percentage_PCA1<-round((eval$V1[1] / sum(eval$V1) )*100 ,2)
percentage_PCA2<-round((eval$V1[2] / sum(eval$V1) )*100 ,2)

plot_1<-ggplot(evec) + 
  geom_point(aes(V3, V4, shape=factor(rep(c("Holstein","Angus"),c(10,12))), color=factor(c("NP","NP","NP","NP","NP","P","P","P","P","P","NP","NP","NP","NP","NP","NP","NP","P","P","P","P","P"))), size=3)+
  scale_shape_manual(values=c("Holstein"= 0,"Angus"=2))+
  scale_color_manual(values=c("P"="#0066b9", "NP"= "#ff7176"))+
  labs(x = paste("PC1: ",percentage_PCA1,"% variance", sep=""), y = paste("PC2: ",percentage_PCA2,"% variance", sep=""))+
  ggtitle("PCA SNPs")+
  theme_minimal(base_size = 15)+
 theme(
    axis.text = element_blank(),
    plot.margin=grid::unit(c(0,0,0,0), "mm"),
    legend.position = "bottom",
    legend.title = element_text(size=10),
    legend.text  = element_text(size=10),
    plot.title = element_text(hjust = 0.5, size=15)
  )
```

```
vst_DESeq <- DESeq2::vst( DESeq2::DESeqDataSetFromMatrix(countData=count_data_filtered, colData=sample_info, design= ~ preg + breed), blind = TRUE, nsub = 5000)

pcaData<-plotPCA(vst_DESeq, intgroup=c("preg", "breed"), returnData=TRUE)
percentVar <- round(100 * attr(pcaData, "percentVar"))

plot_2<-ggplot(pcaData, aes(PC1, PC2, color=preg, shape= breed)) +
  scale_shape_manual(name=NULL, values=c("HOL"= 2,"ANG"=0), labels=c("Holsein", "Angus"))+
  scale_color_manual(name=NULL, values=c("P" = "#0066b9","NP"= "#ff7176"), labels=c("Fertile", "Sub-fertile"))+
  geom_point(size=3) +
  xlab(paste0("PC1: ",percentVar[1],"% variance")) +
  ylab(paste0("PC2: ",percentVar[2],"% variance")) + 
  ggtitle("PCA transcriptome data")+
  theme_minimal(base_size = 15)+
 theme(
    axis.text = element_blank(),
    plot.margin=grid::unit(c(0,0,0,0), "mm"),
    legend.position = "bottom",
    legend.title = element_text(size=10),
    legend.text  = element_text(size=10),
    plot.title = element_text(hjust = 0.5, size=15)
  )
legend<-get_legend( plot_2 )
```

```
colnames(proteome_data)<-rep(heifer_data$SNP_array_ID, each=2)
proteome_data_pca<-proteome_data
proteome_data_pca$proteins<-rownames(proteome_data_pca)
proteome_data_pca<-reshape2::melt(proteome_data_pca)
proteome_data_pca<-data.frame(proteome_data_pca %>% group_by(proteins, variable) %>% dplyr::summarize(Mean = mean(value, na.rm=TRUE)), stringsAsFactors = FALSE)
proteome_data_pca$variable<-as.character(proteome_data_pca$variable)
proteome_data_pca<-proteome_data_pca[!duplicated(proteome_data_pca[,2:3]),]
proteome_data_pca<-reshape2::dcast(proteome_data_pca, proteins ~ variable)

pca_proteome<- prcomp(na.omit(proteome_data_pca[,c(2:23)]))

summary_PCA<-summary(pca_proteome)
PCA1<-round(summary_PCA$importance[2,1]*100,1)
PCA2<-round(summary_PCA$importance[2,2]*100,1)

pca_proteome<-cbind(pca_proteome$rotation,heifer_data)

plot_3<-ggplot(pca_proteome, aes(x=PC1, y=PC2, color=preg, shape= breed)) +
  scale_shape_manual(name=NULL, values=c("HOL"= 2,"ANG"=0), labels=c("Holsein", "Angus"))+
  scale_color_manual(name=NULL, values=c("P" = "#0066b9","NP"= "#ff7176"), labels=c("Fertile", "Sub-fertile"))+
  geom_point(size=3) +
  xlab(paste0("PC1: ",PCA1,"% variance")) +
  ylab(paste0("PC2: ",PCA2,"% variance")) + 
  ggtitle("PCA proteome data")+
  theme_minimal(base_size = 15)+
 theme(
    axis.text = element_blank(),
    plot.margin=grid::unit(c(0,0,0,0), "mm"),
    legend.position = "bottom",
    legend.title = element_text(size=10),
    legend.text  = element_text(size=10),
    plot.title = element_text(hjust = 0.5, size=15)
  )
```

```
fig <- ggdraw() + draw_image(magick::image_read_pdf("/mnt/storage/lab_folder/shared_R_codes/fernando/angus_holstein_association/Fig1_A_B.pdf", density = 600))
cowplot::plot_grid(
  cowplot::plot_grid(fig),
  NULL,
 cowplot::plot_grid(cowplot::plot_grid(plot_1+ theme(legend.position="none"), NULL, plot_2+ theme(legend.position="none"), NULL, plot_3+ theme(legend.position="none") , rel_widths = c(1,0.1,1,0.1,1), nrow=1, labels=c("C", "", "D","","E"),label_fontface = "plain", label_size=15),
                   legend, nrow=2, rel_heights = c(1,0.1)),
                   rel_heights = c(0.8,0.1,1), nrow=3)
```

##### Figure 2

```
angus_holstein_genotypes_qc_data.fisher$CHR<-paste("chr",angus_holstein_genotypes_qc_data.fisher$CHR, sep="")
angus_holstein_genotypes_qc_data.fisher.GRanges<-GenomicRanges::makeGRangesFromDataFrame(angus_holstein_genotypes_qc_data.fisher, ignore.strand=TRUE,seqinfo=NULL,seqnames.field=c("CHR"),  start.field="BP", end.field=c("BP"),keep.extra.columns=TRUE)
ch<-rtracklayer::import.chain("/home/fbiase/genome/lifover_chains/bosTau8ToBosTau9.over.chain")
seqlevelsStyle(angus_holstein_genotypes_qc_data.fisher.GRanges) <-"UCSC"
angus_holstein_genotypes_qc_data.fisher.GRanges_btau9 <- rtracklayer::liftOver(angus_holstein_genotypes_qc_data.fisher.GRanges, ch)
#angus_holstein_genotypes_qc_data.fisher.GRanges_btau9
angus_holstein_genotypes_qc_data.fisher.GRanges_btau9 <- unlist(angus_holstein_genotypes_qc_data.fisher.GRanges_btau9)
genome(angus_holstein_genotypes_qc_data.fisher.GRanges_btau9) <- "btau9"
#angus_holstein_genotypes_qc_data.fisher.GRanges_btau9
angus_holstein_genotypes_qc_data.fisher.GRanges_btau9_df<- data.frame(iranges = angus_holstein_genotypes_qc_data.fisher.GRanges_btau9) 
#write.table(angus_holstein_genotypes_qc_data.fisher.GRanges_btau9_df, "/mnt/storage/lab_folder/shared_R_codes/fernando/angus_holstein_association/results/Additional_file_2.txt",quote = FALSE, sep = "\t",row.names = FALSE, col.names = TRUE)
angus_holstein_genotypes_qc_data.fisher.GRanges_btau9_df$label<-""
angus_holstein_genotypes_qc_data.fisher.GRanges_btau9_df$label<-ifelse(angus_holstein_genotypes_qc_data.fisher.GRanges_btau9_df$iranges.P <= 2.658e-05, paste(angus_holstein_genotypes_qc_data.fisher.GRanges_btau9_df$iranges.seqnames,":", angus_holstein_genotypes_qc_data.fisher.GRanges_btau9_df$iranges.start, sep="") ,"")

plot_3<-ggmanh::manhattan_plot(x = angus_holstein_genotypes_qc_data.fisher.GRanges_btau9_df, pval.colname = "iranges.P", chr.colname = "iranges.seqnames", pos.colname = "iranges.start",  rescale = FALSE, signif = c( 1e-05), label.font.size = 5 , force = 20, chr.order =c("chr1", "chr2", "chr3","chr4","chr5" , "chr6" , "chr7",  "chr8" , "chr9", "chr10", "chr11", "chr12", "chr13", "chr14", "chr15", "chr16", "chr17", "chr18", "chr19", "chr20", "chr21", "chr22", "chr23" ,"chr24", "chr25" ,"chr26" ,"chr27", "chr28", "chr29" ),label.colname ="label")
```

```
ah_genotypes<- data.table::fread("/mnt/storage/lab_folder/heifer_infertility/AI_angus_holstein/assoc_analysis/Virginia_Tech_Univ_Biase_BOV770V01_20220825_FinalReport.txt", header = TRUE, sep = "\t", skip = 9)
ah_genotypes<-ah_genotypes[,c(1,2,3,4)]
colnames(ah_genotypes)<-c("SNP_Name", "Sample_ID","Allele1", "Allele2")
ah_genotypes<-ah_genotypes[ah_genotypes$SNP_Name %in% c("BovineHD1200026258","Hapmap31720-BTA-126418"),]
ah_genotypes$Sample_ID <- gsub(" ","_",ah_genotypes$Sample_ID)
ah_genotypes<- ah_genotypes[!ah_genotypes$Sample_ID=="Sample_23" & !ah_genotypes$Sample_ID=="Sample_24",]
ah_genotypes$genotype<-paste(ah_genotypes$Allele1,ah_genotypes$Allele2, sep="")
#ah_genotypes %>% group_by(SNP_Name, genotype) %>% tally()
sample_id <- c("Sample_1","Sample_2","Sample_3","Sample_4","Sample_5","Sample_6","Sample_7","Sample_8","Sample_9","Sample_10","Sample_11","Sample_12","Sample_13","Sample_14","Sample_15","Sample_16","Sample_17","Sample_18","Sample_19","Sample_20","Sample_21","Sample_22") 
breed<-c("hosltein","hosltein","hosltein","hosltein","hosltein","hosltein","hosltein","hosltein","hosltein","hosltein","angus","angus","angus","angus","angus","angus","angus","angus","angus","angus","angus","angus")
phenotype_angus_holstein <- c("2","2","2","2","2","1","1","1","1","1","2","2","2","2","2", "2","2","1","1","1","1","1")
ah_genotypes<-merge(ah_genotypes,data.frame(sample_id,phenotype_angus_holstein, breed), by.x="Sample_ID", by.y="sample_id")
ah_genotypes<-ah_genotypes[!(ah_genotypes$genotype == "--"),]
ah_genotypes_count<-ah_genotypes %>% group_by(SNP_Name, genotype,phenotype_angus_holstein) %>% tally() %>% arrange(phenotype_angus_holstein, .by_group = TRUE )

supp.labs <- c("rs110918927", "rs109366560")
names(supp.labs) <- c("BovineHD1200026258", "Hapmap31720-BTA-126418")

plot_4<-ggplot(ah_genotypes_count, aes(fill= genotype , y=n, x=phenotype_angus_holstein))+
geom_bar(position="fill", stat="identity")+
scale_fill_manual(values=c("AA" = "#cb6054","AG"= "#87a14d", "GG"="#9c72be"))+
scale_y_continuous(name="Frequency")+
scale_x_discrete(name=NULL, labels=c("Fertile", "Sub-fertile"))+
facet_grid(~SNP_Name, labeller = labeller(SNP_Name = supp.labs))+
    theme_classic(base_size=15)+
    theme(
    axis.text.x=element_text(color="black", angle=45, hjust=1),
    strip.background = element_blank(),
    legend.title=element_blank(),
    legend.position="bottom"
)
```

```
plot_grid(plot_3,plot_4, nrow=1, rel_widths=c(1,0.25))
```

##### Figure 3

```
plotdata <- data_cpm_filtered
plotdata <- plotdata[rownames(plotdata)=="ENSBTAG00000004278",]
plotdata <- as.data.frame(plotdata)
plotdata <- cbind(heifer_data, plotdata)
plotdata$shape <- c(0:9, 0:11)

plotdata2 <- data_cpm_filtered
plotdata2 <- plotdata2[rownames(plotdata2)=="ENSBTAG00000002972",]
plotdata2 <- as.data.frame(plotdata2)
plotdata2 <- cbind(heifer_data, plotdata2)
plotdata2$shape <- c(0:9, 0:11)
```

```
plotdata %>% group_by(breed, preg) %>% dplyr::summarize(Mean = mean(plotdata, na.rm=TRUE))
```

```
## `summarise()` has grouped output by 'breed'. You can override using the
## `.groups` argument.
```

```
## # A tibble: 4 × 3
## # Groups:   breed [2]
##   breed preg   Mean
##   <chr> <chr> <dbl>
## 1 ANG   NP     49.9
## 2 ANG   P      55.1
## 3 HOL   NP     40.7
## 4 HOL   P      50.1
```

```
plotdata2 %>% group_by(breed, preg) %>% dplyr::summarize(Mean = mean(plotdata2, na.rm=TRUE))
```

```
## `summarise()` has grouped output by 'breed'. You can override using the
## `.groups` argument.
```

```
## # A tibble: 4 × 3
## # Groups:   breed [2]
##   breed preg   Mean
##   <chr> <chr> <dbl>
## 1 ANG   NP     4.44
## 2 ANG   P      3.64
## 3 HOL   NP     5.85
## 4 HOL   P      4.73
```

```
font_size=12
plot_5 <- ggplot(data=plotdata, aes(y=plotdata, x=breed, color=preg))+
geom_point(position=position_jitterdodge(jitter.width = 0.5,jitter.height = 0,dodge.width = 1),size=2, shape=plotdata$shape)+
  scale_color_manual(labels=c("Sub-fertile", "Fertile"), values=c("#ff7176", "#0066b9"))+
  scale_x_discrete(name =NULL, labels=c("Angus", "Holstein"))+
  ylab("CPM")+
  ggtitle("APMAP\nadipocyte plasma membrane associated protein")+
  theme_classic2()+
  theme(legend.key = element_rect(),
        legend.position = "none",
        plot.title = element_text(size=font_size, face = "italic"),
        axis.text.x = element_text(size=font_size, color="black"),
        axis.text.y = element_text(size=font_size, color="black"),
        axis.title.y = element_text(size=font_size, color="black"),
        axis.title.x = element_text(size=font_size, color="black"))

plot_6 <- ggplot(data=plotdata2, aes(y=plotdata2, x=breed, color=preg))+
geom_point(position=position_jitterdodge(jitter.width = 0.5,jitter.height = 0,dodge.width = 1),size=2, shape=plotdata$shape)+
  scale_color_manual(labels=c("Sub-fertile", "Fertile"), values=c("#ff7176", "#0066b9"))+
  scale_x_discrete(name =NULL, labels=c("Angus", "Holstein"))+
  ylab("")+
  ggtitle("DNAI7\ndynein axonemal intermediate chain 7")+
  theme_classic2()+
  theme(legend.key = element_rect(),
        legend.position = "bottom",
        legend.title = element_blank(),
        legend.text = element_text(size=font_size, color="black"),
        plot.title = element_text(size=font_size, face = "italic"),
        axis.text.x = element_text(size=font_size, color="black"),
        axis.text.y = element_text(size=font_size, color="black"),
        axis.title.y = element_text(size=font_size, color="black"),
        axis.title.x = element_text(size=font_size, color="black"))
legend <- get_legend(plot_6)


plots <- plot_grid(plot_5, plot_6+theme(legend.position="none"), nrow=1)
plot_grid(plots, legend, nrow=2, ncol=1, rel_heights = c(1, .1))
```

##### Figure 4

```
colnames(proteome_data)<-c("07","07","08","08","09","09","10","10","11","11","12","12","13","13","14","14","15","15","16","16","17","17","18","18","19","19","20","20","21","21","22","22","23","23","24","24","25","25","26","26","27","27","28","28")
heifer_data <- sample_info

plotdata <- proteome_data
plotdata <- plotdata[rownames(plotdata)=="A5D798",]
plotdata <- reshape2::melt(plotdata)
```

```
## No id variables; using all as measure variables
```

```
plotdata <- merge(heifer_data, plotdata, by.x="sample", by.y="variable")

plotdata2 <- proteome_data
plotdata2 <- plotdata2[rownames(plotdata2)=="P19034",]
plotdata2 <- reshape2::melt(plotdata2)
```

```
## No id variables; using all as measure variables
```

```
plotdata2 <- merge(heifer_data, plotdata2, by.x="sample", by.y="variable")

plotdata3 <- proteome_data
plotdata3 <- plotdata3[rownames(plotdata3)=="F1MYX5",]
plotdata3 <- reshape2::melt(plotdata3)
```

```
## No id variables; using all as measure variables
```

```
plotdata3 <- merge(heifer_data, plotdata3, by.x="sample", by.y="variable")

plotdata<- plotdata %>% group_by(breed, preg,Animal_ID) %>% dplyr::summarize(Mean = mean(value, na.rm=TRUE))
```

```
## `summarise()` has grouped output by 'breed', 'preg'. You can override using the
## `.groups` argument.
```

```
plotdata2<-plotdata2 %>% group_by(breed, preg,Animal_ID) %>% dplyr::summarize(Mean = mean(value, na.rm=TRUE))
```

```
## `summarise()` has grouped output by 'breed', 'preg'. You can override using the
## `.groups` argument.
```

```
plotdata3<-plotdata3 %>% group_by(breed, preg,Animal_ID) %>% dplyr::summarize(Mean = mean(value, na.rm=TRUE))
```

```
## `summarise()` has grouped output by 'breed', 'preg'. You can override using the
## `.groups` argument.
```

```
plotdata$shape <- c(c(0,1,2,3,4,5,6,7,8,9),
                    c(0,1,2,3,4,5,6,7,8,9,10,11))

plotdata2$shape <- c(c(0,1,2,3,4,5,6,7,8,9),
                    c(0,1,2,3,4,5,6,7,8,9,10,11))

plotdata3$shape <- c(c(0,1,2,3,4,5,6,7,8,9),
                    c(0,1,2,3,4,5,6,7,8,9,10,11))

font_size=12

plot_7 <- ggplot(data=plotdata, aes(y=Mean, x=breed, color=preg))+
  geom_point(position=position_jitterdodge(jitter.width = 0.5,jitter.height = 0,dodge.width = 1),size=3, shape=plotdata$shape)+
  scale_color_manual(labels=c("Sub-fertile", "Fertile"), values=c("#ff7176", "#0066b9"))+
  scale_x_discrete(name =NULL, labels=c("Angus", "Holstein"))+
  ylab("Log(Protein abundance)")+
  ggtitle("Alpha-ketoglutarate-dependent dioxygenase FTO")+
  theme_classic2()+
  annotate("text", x = c(0.9,2), y = c(32.5,32), label = c("estimate = -0.395\nP = 0.0119","estimate = -0.747\nP = 0.0001"))+
  theme(legend.key = element_rect(),
        legend.position = "none",
        plot.title = element_text(size=font_size, face = "italic"),
        axis.text.x = element_text(size=font_size, color="black"),
        axis.text.y = element_text(size=font_size, color="black"),
        axis.title.y = element_text(size=font_size, color="black"),
        axis.title.x = element_text(size=font_size, color="black"))

plot_8 <- ggplot(data=plotdata3, aes(Mean, x=breed, color=preg))+
  geom_point(position=position_jitterdodge(jitter.width = 0.5,jitter.height = 0,dodge.width = 1),size=3, shape=plotdata$shape)+
  scale_color_manual(labels=c("Sub-fertile", "Fertile"), values=c("#ff7176", "#0066b9"))+
  scale_x_discrete(name =NULL, labels=c("Angus", "Holstein"))+
  ylab("")+
  ggtitle("Apolipoprotein C-II")+
  annotate("text", x = c(0.9,2), y = c(24,26), label = c("estimate = 0.546\nP = 0.1757","estimate = 2.325\nP = <0.0001"))+
  theme_classic2()+
  theme(legend.key = element_rect(),
        legend.position = "none",
        legend.title = element_blank(),
        legend.text = element_text(size=font_size, color="black"),
        plot.title = element_text(size=font_size, face = "italic"),
        axis.text.x = element_text(size=font_size, color="black"),
        axis.text.y = element_text(size=font_size, color="black"),
        axis.title.y = element_text(size=font_size, color="black"),
        axis.title.x = element_text(size=font_size, color="black"))

plot_9 <- ggplot(data=plotdata2, aes(Mean, x=breed, color=preg))+
  geom_point(position=position_jitterdodge(jitter.width = 0.5,jitter.height = 0,dodge.width = 1),size=3, shape=plotdata$shape)+
  scale_color_manual(labels=c("Sub-fertile", "Fertile"), values=c("#ff7176", "#0066b9"))+
  scale_x_discrete(name =NULL, labels=c("Angus", "Holstein"))+
  ylab("")+
  ggtitle("Lymphocyte cytosolic protein 1 ")+
  annotate("text", x = c(0.9,2), y = c(26,28), label = c("estimate = 0.429\nP = 0.2202","estimate = 1.855\nP = 0.0001"))+
  theme_classic2()+
  theme(legend.key = element_rect(),
        legend.position = "bottom",
        legend.title = element_blank(),
        legend.text = element_text(size=font_size, color="black"),
        plot.title = element_text(size=font_size, face = "italic"),
        axis.text.x = element_text(size=font_size, color="black"),
        axis.text.y = element_text(size=font_size, color="black"),
        axis.title.y = element_text(size=font_size, color="black"),
        axis.title.x = element_text(size=font_size, color="black"))
   
legend <- get_legend(plot_9)

plots <- plot_grid(plot_7,plot_8, plot_9+theme(legend.position = "none",), nrow=1, labels=c("A", "B", "C"),label_fontface = "plain",label_size = 13)
plot_grid(plots, legend, nrow=2, ncol=1, rel_heights = c(1, .1))
```

##### Figure 5

```
plot_10<-plot_variance_explained(MOFAobject.trained.group, x="view", y="factor")+
theme(axis.text.x=element_text(size=10,angle=45,vjust=0.5,hjust=0.5),
      axis.text.y=element_text(size=10),
      plot.title=element_text(size=10),
      strip.text = element_text(size = 10),
      legend.position="top",
      plot.margin=unit(x=c(0,0,0,0),units="mm"),
      legend.margin=margin(c(0,0,0,0), unit='mm'))


plot_11<-plot_factor(MOFAobject.trained.group, 
  factor = 1:3,
  color_by = "pregnancy",
  shape_by = "breed"
) +
scale_fill_manual(name=NULL, values=c("Fertile" = "#0066b9","Sub-fertile"= "#ff7176"), labels=c("Fertile", "Sub-fertile"))+
scale_shape_manual(name=NULL, values=c(21,23))+
theme(axis.text.y=element_text(size=10),
      axis.text.x=element_text(size=10,angle=45,vjust=0.5,hjust=0.5),
      strip.text = element_text(size = 10),
      legend.position="bottom",
      legend.spacing.x = unit(0, 'cm'))
```

```
## Scale for shape is already present.
## Adding another scale for shape, which will replace the existing scale.
```

```
body(plot_top_weights)[[22]][[3]][[2]][[3]][[3]][[4]]<-1

plot_12<-plot_top_weights(MOFAobject.trained.group,
  view = "genotypes",
  factor = c(1:3),
  nfeatures = 10)+
  theme(
axis.title.x=element_blank(),
axis.text.y = element_text(size = 8),
strip.text = element_text(size = 10)
)

plot_13<-plot_top_weights(MOFAobject.trained.group,
  view = "mRNA",
  factor = c(1:3),
  nfeatures = 10)+
  theme(
axis.title.x=element_blank(),
axis.text.y = element_text(size = 8),
strip.text = element_text(size = 10)
)

plot_14<-plot_top_weights(MOFAobject.trained.group,
  view = "protein",
  factor = c(1:3),
  nfeatures = 10)+
  theme(
axis.text.y = element_text(size = 8),
strip.text = element_text(size = 10),
plot.margin=unit(x=c(t = 0, r = 2, b = 0, l = 20),units="mm"),
panel.spacing = unit(c(5,3), "lines")
)


MOFAobject.trained.group <- run_tsne(MOFAobject.trained.group,perplexity = 5)

plot_15<-plot_dimred(MOFAobject.trained.group,
  method = "TSNE",  # method can be either "TSNE" or "UMAP"
  dot_size=3,
  color_by = "pregnancy",
  shape_by = "breed"
)+
scale_fill_manual(values=c("Fertile" = "#0066b9","Sub-fertile"= "#ff7176"), labels=c("Fertile", "Sub-fertile"))+
ggtitle("t-Distributed Stochastic \n Neighbor Embedding")+
theme(legend.position="none",
plot.title = element_text(size=10, hjust=0.5))
```

```
## Warning: The `<scale>` argument of `guides()` cannot be `FALSE`. Use "none" instead as
## of ggplot2 3.3.4.
## ℹ The deprecated feature was likely used in the MOFA2 package.
##   Please report the issue at <]8;;https://github.com/bioFAM/MOFA2https://github.com/bioFAM/MOFA2]8;;>.
```

```
cowplot::plot_grid(
cowplot::plot_grid(plot_10,plot_11, plot_15,nrow=3, labels = c("A","B","D"),label_size = 13, label_fontface = "plain"),
cowplot::plot_grid(cowplot::plot_grid(plot_12,plot_13, plot_14, nrow=3, rel_heights=c(0.9,0.9,1)),labels = c("C"),label_size = 13, label_fontface = "plain"), 
ncol=2, rel_widths=c(0.4,1))
```

cowplot::plot\_grid(plot\_10,plot\_11, plot\_15,nrow=1, labels =
c(“A”,“B”,“C”),label\_size = 13, label\_fontface = “plain”)

##### plot eFDR

```
efdr<-read.delim("/mnt/storage/lab_folder/shared_R_codes/fernando/angus_holstein_association/results/2022_10_09_efdr.txt")
font_size<-9

plot1<-ggplot()+
geom_point(data=efdr, aes(x=e.pvalue.edgeR , y=raw.pvalue),color="black", size=2,shape=16)+
geom_line(data=efdr, aes(x=e.pvalue.edgeR , y=raw.pvalue),color="black", size=0.1,linetype=3)+
scale_x_continuous(name="empirical FDR", limits = c(0, 0.0005), breaks=seq(0, 0.0005, 0.0001))+
scale_y_continuous(name="nominal P value", limits = c(0, 0.002), breaks=seq(0, 0.002, 0.0005))+
ggtitle("Differential gene expression eFDR \n edgeR")+
theme_bw()+
theme(panel.grid= element_blank(),
panel.background = element_blank(),
panel.grid.minor = element_blank(),
panel.grid.major = element_line(color="lightgray"),
plot.background = element_blank(),
axis.text.y=element_text(color="black", size=font_size),
axis.text.x=element_text(color="black", size=font_size, angle=90),
panel.spacing = unit(c(0.4,0.4,0.4,0.4),"cm"),
plot.margin = unit(c(0.5,0.5,0.5,0.5),"cm"),
legend.position="none",
plot.title = element_text(lineheight=.8, hjust=0.5, size= font_size))

plot2<-ggplot()+
geom_point(data=efdr, aes(x=e.pvalue.DESEQ2 , y=raw.pvalue.1),color="black", size=2,shape=16)+
geom_line(data=efdr, aes(x=e.pvalue.DESEQ2 , y=raw.pvalue.1),color="black", size=0.1,linetype=3)+
scale_x_continuous(name="empirical FDR", limits = c(0, 0.005), breaks=seq(0, 0.05, 0.0005))+
scale_y_continuous(name="nominal P value", limits = c(0, 0.002), breaks=seq(0, 0.002, 0.0005))+
ggtitle("Differential gene expression eFDR \n DeSeq2 WALD test")+
theme_bw()+
theme(panel.grid= element_blank(),
panel.background = element_blank(),
panel.grid.minor = element_blank(),
panel.grid.major = element_line(color="lightgray"),
plot.background = element_blank(),
axis.text.y=element_text(color="black", size=font_size),
axis.text.x=element_text(color="black", size=font_size, angle=90),
panel.spacing = unit(c(0.4,0.4,0.4,0.4),"cm"),
plot.margin = unit(c(0.5,0.5,0.5,0.5),"cm"),
legend.position="none",
plot.title = element_text(lineheight=.8, hjust=0.5, size= font_size))

plot3<-ggplot()+
geom_point(data=efdr, aes(x=e.pvalue.DESEQ2.1 , y=raw.pvalue.1),color="black", size=2,shape=16)+
geom_line(data=efdr, aes(x=e.pvalue.DESEQ2.1 , y=raw.pvalue.1),color="black", size=0.1,linetype=3)+
scale_x_continuous(name="empirical FDR", limits = c(0, 0.005), breaks=seq(0, 0.05, 0.0005))+
scale_y_continuous(name="nominal P value", limits = c(0, 0.002), breaks=seq(0, 0.002, 0.0005))+
ggtitle("Differential gene expression eFDR \n LTR test")+
theme_bw()+
theme(panel.grid= element_blank(),
panel.background = element_blank(),
panel.grid.minor = element_blank(),
panel.grid.major = element_line(color="lightgray"),
plot.background = element_blank(),
axis.text.y=element_text(color="black", size=font_size),
axis.text.x=element_text(color="black", size=font_size, angle=90),
panel.spacing = unit(c(0.4,0.4,0.4,0.4),"cm"),
plot.margin = unit(c(0.5,0.5,0.5,0.5),"cm"),
legend.position="none",
plot.title = element_text(lineheight=.8, hjust=0.5, size= font_size))


cowplot::plot_grid(plot1, plot2, plot3, nrow=1)
```

#### sessionInfo

```
sessionInfo()
```

```
## R version 4.2.2 Patched (2022-11-10 r83330)
## Platform: x86_64-pc-linux-gnu (64-bit)
## Running under: Ubuntu 20.04.5 LTS
## 
## Matrix products: default
## BLAS:   /usr/lib/x86_64-linux-gnu/blas/libblas.so.3.9.0
## LAPACK: /usr/lib/x86_64-linux-gnu/lapack/liblapack.so.3.9.0
## 
## locale:
##  [1] LC_CTYPE=en_US.UTF-8       LC_NUMERIC=C              
##  [3] LC_TIME=en_US.UTF-8        LC_COLLATE=en_US.UTF-8    
##  [5] LC_MONETARY=en_US.UTF-8    LC_MESSAGES=en_US.UTF-8   
##  [7] LC_PAPER=en_US.UTF-8       LC_NAME=C                 
##  [9] LC_ADDRESS=C               LC_TELEPHONE=C            
## [11] LC_MEASUREMENT=en_US.UTF-8 LC_IDENTIFICATION=C       
## 
## attached base packages:
##  [1] parallel  grid      stats4    stats     graphics  grDevices utils    
##  [8] datasets  methods   base     
## 
## other attached packages:
##  [1] MOFA2_1.6.0                 MultiAssayExperiment_1.22.0
##  [3] emmeans_1.8.2               lme4_1.1-31                
##  [5] Matrix_1.5-3                ggmanh_1.0.0               
##  [7] doParallel_1.0.17           iterators_1.0.14           
##  [9] foreach_1.5.2               bigmemory_4.6.1            
## [11] gridExtra_2.3               kableExtra_1.3.4           
## [13] ggsignif_0.6.4              data.table_1.14.6          
## [15] goseq_1.48.0                geneLenDataBase_1.32.0     
## [17] BiasedUrn_2.0.8             car_3.1-1                  
## [19] carData_3.0-5               ggpubr_0.5.0               
## [21] reshape2_1.4.4              htmlwidgets_1.5.4          
## [23] forcats_0.5.2               stringr_1.5.0              
## [25] purrr_0.3.5                 readr_2.1.3                
## [27] tidyr_1.2.1                 tibble_3.1.8               
## [29] tidyverse_1.3.2             plotly_4.10.1              
## [31] flashClust_1.01-2           ComplexHeatmap_2.12.1      
## [33] VennDiagram_1.7.3           futile.logger_1.4.3        
## [35] readxl_1.4.1                DESeq2_1.36.0              
## [37] SummarizedExperiment_1.26.1 Biobase_2.56.0             
## [39] MatrixGenerics_1.8.0        matrixStats_0.63.0         
## [41] GenomicRanges_1.48.0        GenomeInfoDb_1.32.2        
## [43] IRanges_2.30.0              S4Vectors_0.34.0           
## [45] BiocGenerics_0.42.0         GGally_2.1.2               
## [47] cowplot_1.1.1               edgeR_3.38.4               
## [49] limma_3.52.1                ggplot2_3.4.0              
## [51] dplyr_1.0.10               
## 
## loaded via a namespace (and not attached):
##   [1] estimability_1.4.1       rappdirs_0.3.3           rtracklayer_1.56.0      
##   [4] coda_0.19-4              bit64_4.0.5              knitr_1.41              
##   [7] multcomp_1.4-20          DelayedArray_0.22.0      KEGGREST_1.36.3         
##  [10] RCurl_1.98-1.7           generics_0.1.3           GenomicFeatures_1.48.3  
##  [13] lambda.r_1.2.4           TH.data_1.1-1            RSQLite_2.2.19          
##  [16] bit_4.0.5                tzdb_0.3.0               webshot_0.5.4           
##  [19] xml2_1.3.3               lubridate_1.9.0          assertthat_0.2.1        
##  [22] gargle_1.2.1             xfun_0.35                hms_1.1.2               
##  [25] jquerylib_0.1.4          evaluate_0.18            fansi_1.0.3             
##  [28] restfulr_0.0.14          progress_1.2.2           dbplyr_2.2.1            
##  [31] DBI_1.1.3                geneplotter_1.74.0       reshape_0.8.9           
##  [34] googledrive_2.0.0        ellipsis_0.3.2           corrplot_0.92           
##  [37] backports_1.4.1          annotate_1.74.0          biomaRt_2.52.0          
##  [40] vctrs_0.5.1              abind_1.4-5              cachem_1.0.6            
##  [43] withr_2.5.0              GenomicAlignments_1.32.0 prettyunits_1.1.1       
##  [46] svglite_2.1.0            cluster_2.1.4            dir.expiry_1.4.0        
##  [49] lazyeval_0.2.2           crayon_1.5.2             basilisk.utils_1.8.0    
##  [52] genefilter_1.78.0        labeling_0.4.2           pkgconfig_2.0.3         
##  [55] nlme_3.1-160             rlang_1.0.6              lifecycle_1.0.3         
##  [58] sandwich_3.0-2           bigmemory.sri_0.1.6      filelock_1.0.2          
##  [61] BiocFileCache_2.4.0      modelr_0.1.10            cellranger_1.1.0        
##  [64] Rhdf5lib_1.18.2          boot_1.3-28.1            zoo_1.8-11              
##  [67] reprex_2.0.2             GlobalOptions_0.1.2      googlesheets4_1.0.1     
##  [70] pheatmap_1.0.12          png_0.1-8                viridisLite_0.4.1       
##  [73] rjson_0.2.21             bitops_1.0-7             rhdf5filters_1.8.0      
##  [76] Biostrings_2.64.1        blob_1.2.3               shape_1.4.6             
##  [79] pdftools_3.2.1           qpdf_1.2.0               rstatix_0.7.1           
##  [82] scales_1.2.1             memoise_2.0.1            magrittr_2.0.3          
##  [85] plyr_1.8.8               zlibbioc_1.42.0          compiler_4.2.2          
##  [88] BiocIO_1.6.0             RColorBrewer_1.1-3       clue_0.3-63             
##  [91] Rsamtools_2.12.0         cli_3.4.1                XVector_0.36.0          
##  [94] formatR_1.12             MASS_7.3-58.1            mgcv_1.8-41             
##  [97] tidyselect_1.2.0         stringi_1.7.8            highr_0.9               
## [100] yaml_2.3.6               askpass_1.1              locfit_1.5-9.6          
## [103] ggrepel_0.9.2            sass_0.4.4               tools_4.2.2             
## [106] timechange_0.1.1         circlize_0.4.15          rstudioapi_0.14         
## [109] uuid_1.1-0               farver_2.1.1             Rtsne_0.16              
## [112] digest_0.6.30            Rcpp_1.0.9               broom_1.0.1             
## [115] httr_1.4.4               AnnotationDbi_1.58.0     colorspace_2.0-3        
## [118] rvest_1.0.3              XML_3.99-0.13            fs_1.5.2                
## [121] reticulate_1.26          splines_4.2.2            statmod_1.4.37          
## [124] uwot_0.1.14              basilisk_1.8.1           systemfonts_1.0.4       
## [127] xtable_1.8-4             jsonlite_1.8.4           nloptr_2.0.3            
## [130] futile.options_1.0.1     R6_2.5.1                 pillar_1.8.1            
## [133] htmltools_0.5.3          mime_0.12                glue_1.6.2              
## [136] fastmap_1.1.0            minqa_1.2.5              BiocParallel_1.30.4     
## [139] codetools_0.2-18         mvtnorm_1.1-3            utf8_1.2.2              
## [142] lattice_0.20-45          bslib_0.4.1              pbkrtest_0.5.1          
## [145] curl_4.3.3               magick_2.7.3             GO.db_3.15.0            
## [148] survival_3.4-0           rmarkdown_2.17           munsell_0.5.0           
## [151] GetoptLong_1.0.5         rhdf5_2.40.0             GenomeInfoDbData_1.2.8  
## [154] HDF5Array_1.24.2         haven_2.5.1              gtable_0.3.1
```
